# Neuronal lysosome transfer to oligodendrocyte precursor cells: a novel mechanism of neuron-glia communication and its role in neurodegenerative diseases

**DOI:** 10.1101/2024.03.03.583173

**Authors:** Li-Pao Fang, Ching-Hsin Lin, Yasser Medlej, Renping Zhao, Hsin-Fang Chang, Yixun Su, Na Zhao, Davide Gobbo, Qilin Guo, Amanda Wyatt, Vanessa Wahl, Ulrich Boehm, Wenhui Huang, Marcel A. Lauterbach, Chenju Yi, Jianqin Niu, Anja Scheller, Frank Kirchhoff, Xianshu Bai

**Author notes:** Present address: Institute of Anatomy and Cell Biology, University of Saarland, 66421 Homburg, Germany (Na Zhao). Correspondence should be addressed to Prof. Frank Kirchhoff, Dr. Xianshu Bai.

## Abstract

Oligodendrocyte precursor cells (OPCs) shape brain function through intricate regulatory mechanisms. Here, we observed that OPC processes establish connections with neuronal somata, with smaller lysosomes positioned near these contact sites. Tracking lysosomes demonstrated neuronal lysosomes were attracted to and released at these contact points, eventually becoming incorporated into OPC processes, suggesting a selective, OPC-evoked release of lysosomes from neuronal soma and their ingestion by OPCs, highlighting a unique lysosome-mediated communication between neurons and OPCs. Diminished branching of OPC processes resulted in fewer neuron-OPC contacts, fostering larger lysosome accumulation in neurons, altered neuronal activity and escalated prevalence of senescent neurons during aging. A similar reduction in OPC branching and neuronal lysosome accumulation was evident in an early-stage Alzheimer’s disease mouse model. Together, these findings underscore the pivotal role of OPC processes in modulating neuronal activity through direct somatic contact and lysosome ingestion, presenting a prospective therapeutic avenue for addressing neurodegenerative diseases.

## Introduction

Oligodendrocyte precursor cells (OPCs) exhibit complex morphology with small cell bodies and branched, motile processes that survey their local microenvironment^1^. Under pathological conditions, including acute brain injury or epilepsy, OPCs often become hypertrophic, elevating their morphological complexity^2^. This phenomenon correlates with an increase of neuronal activity^3, 4^, yet the causative relationship between the changes in OPC morphology and neuronal activity remains unclear.

During development or in response to acute brain injury, OPCs adeptly sense changes in brain activity, transiently suppressing the transcription factor Olig2 and remain in an undifferentiated state, contributing to brain remodeling^5^. OPCs also respond to neuronal activity through the expression of glutamatergic and GABAergic receptors^6, 7^ on post-synaptic terminals^8, 9^. While conventional understanding describes OPCs forming classical post-synapses, recent studies unveil diverse pathways for OPC-neuron interactions, particularly between OPC processes and neurons^10^, including GABA release to neurons in the hippocampus via synaptic complexes^11^, synaptic pruning by phagocytosing axonal terminals during development^12^, and putative modulation of action potential propagation through contacts at the nodes of Ranvier^13^.

The complex morphology of OPCs suggests a mode of communication with the densely populated neurons in the brain, potentially through alternative physical contacts such as direct interactions with neuronal somata.

We, thus, investigated the interaction between OPC processes and neurons employing various transgenic mouse models and advanced imaging techniques. Our findings revealed that OPC processes exhibit equal efficacy in contacting the somata of both inhibitory and excitatory neurons, but favouring active neurons. Notably, smaller lysosomes within the neuronal compartment were located in closer proximity to the contact sites. Live imaging of lysosomes in neuron-OPC co-cultures showed that neuronal lysosomes were recruited to and released at the contact sites, and eventually appeared in OPC processes, implying that OPCs might not only facilitate neuronal lysosome release by forming contacts but also by actively engulfing neuronal lysosomes. Furthermore, in genetically modified mice with reduced branching of OPC processes, due to OPC-specific deletion of L-type calcium channel Cav1.2 and Cav1.3 genes, fewer OPC-neuron contacts, aberrant lysosome accumulation in neuronal soma during adulthood and enhanced senescence of neurons in aging were observed. Similar reductions in OPC-neuron contacts and accumulation of neuronal lysosomes were present in an early-stage Tg2576 Alzheimer’s disease mouse model, linking OPC-neuron contact and neurodegeneration. In conclusion, our results demonstrate a critical role of OPC contacts with neuronal somata for neuronal lysosome release and maintaining proper neuronal activity, underscoring the clinical significance of this mechanism, particularly in the context of neurodegenerative conditions.

## Results

### OPC processes contacted neuronal somata with a preference for active neurons

To explore the possibility of OPCs forming contacts with neurons beyond the synaptic connections formed by neuronal axons on the OPC surface, we analysed the spatial relationship between OPCs and neurons by performing GFP and NeuN double immunostaining in adult NG2-EYFP (NG2^EYFP^) mouse brains (**Fig. 1A**). In the cortex, GFP^+^ OPCs exhibited a ramified morphology, and several processes were found in contact with neuronal somata (**Fig. 1B, C**). Such process-soma junctions were commonly observed for the majority, if not all, of the neurons (ranging from 91.9-99.3%) in the grey matter, including cortex, hippocampus, thalamus, hypothalamus, and amygdala in both developing and adult brains (**Fig. 1D**, **Suppl. Fig. 1, 2**). However, we noticed a variability in the number of contacts among neurons. Therefore, we further compared the contact probability between excitatory and inhibitory neurons by performing triple immunostaining of GFP, NeuN, and GABA in NG2^EYFP^ mice (**Fig. 1E**). A similar frequency of contact was observed between excitatory (NeuN^+^GABA^-^) and inhibitory neurons (NeuN^+^GABA^+^) (**Fig. 1E, F**; **Suppl. Fig. 1C**).

**Figure 1.**
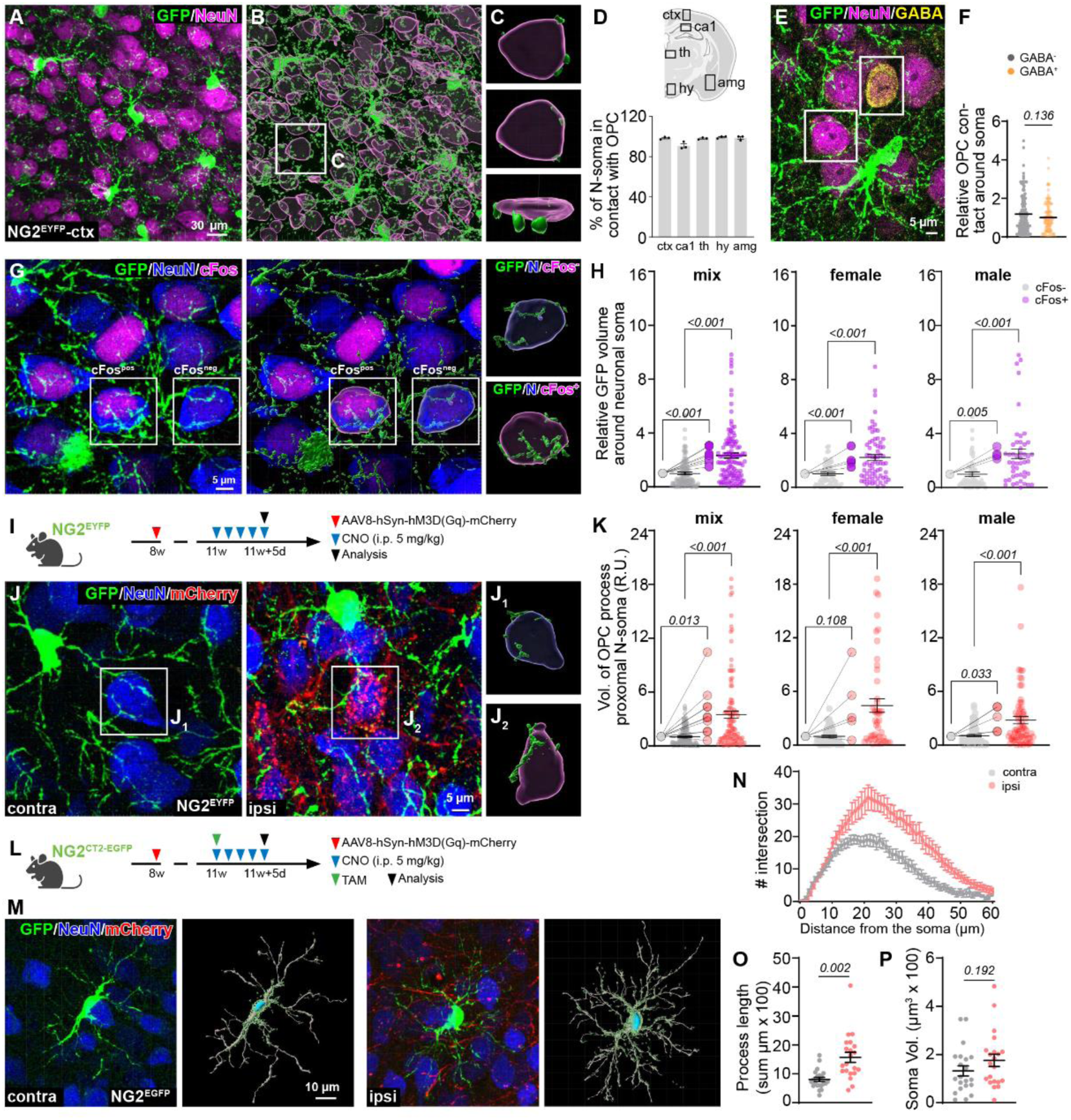
OPC processes contact neuronal somata with a preference for active neurons. A, B. Immunostaining and 3D reconstruction of OPCs and neurons with GFP and NeuN antibodies in coronal brain slices of NG2-EYFP (NG2^EYFP^) mice. **C** Exemplary images of process-somata contact between OPCs and neurons from the boxed area of **B**, shown in three directions. **D** Quantification of OPC-neuron contact frequency for neurons in the cortex (ctx), hippocampal CA1 region (ca1), thalamus (th), hypothalamus (hy) and amygdala (amg). **E, F** Immunostaining and quantification of OPC (GFP^+^) contacts on GABAergic (GABA^+^) and non-GABAergic (GABA^-^) neurons in NG2^EYFP^ mice. **G** Immunostaining of OPCs and neurons with GFP and NeuN in coronal brain slices of NG2^EYFP^ mice. Neuronal activity was indicated with cFos (magenta) immunoreactivity. **H** Quantification of relative volume (Vol.) of OPC processes (GFP^+^) contacting cFos^+^ and cFos^-^ neurons, in both female and male mice. (female: 68 cFos^-^ cells and 70 cFos^+^ cells in N=4 mice; male: 49 cFos^-^ cells and 48 cFos^+^ cells in N=4 mice, two-sided unpaired t-tests for cells and two-sided paired t-tests for mice). **I** Experimental scheme for **J and K**. **J** Immunostaining and 3D reconstruction of OPCs and neurons with GFP and NeuN antibodies in contra-and ipsilateral side. mCherry expression indicates neurons transfected with hSyn-hM3Dq virus that can be activated by CNO application. **K** Quantification of the volume of OPC processes in contact with mCherry^+^ and mCherry^-^ neuronal soma (female: 62 mCherry^-^ cells and 46 mCherry ^+^ cells in N=5 mice; male: 92 mCherry^-^ cells and 65 mCherry^+^ cells in N=5 mice, two-sided unpaired t-tests for cells and two-sided paired t-tests for mice). **L** Experimental scheme for **M-P**. **M** Immunostaining and 3D reconstruction of single OPC with GFP in contra– and ipsilateral side in NG2-CreER^T2^ x CAG-^fl^CTA^fl^-EGFP. **N-P** Analysis of OPC morphology in contra– and ipsilateral side in terms of the number of processes intersection (**N**), total length of processes (**O**) and soma volume (**P**) (contra: n=21 cells, ipsi: n=21 cells; two-sided unpaired t-tests). Scale bars in **A**=30 µm, **E, G, J**=5 µm, **M**=10 µm.

Since OPCs sense neuronal activity^10, 14^, we then further investigated whether such contact is related to neuronal activity. Distinguishing between active and less/non-active neurons based on immunoreactivity to cFos (**Fig. 1G**), an immediate-early gene and a well-established marker of cellular activity^15, 16^, we observed that cFos^+^ neurons exhibited approximately twice as many contacts with OPC processes compared to cFos^-^ neurons in both female and male mice (**Fig. 1G, H**, **Suppl. Fig. 3**). These findings strongly suggested a preference of OPCs for contacting active neurons. To validate this observation, we employed a chemogenetic approach to activate cortical neurons. By injecting AAV8-hSyn-hM3D(Gq)-mCherry virus intracortically and administering CNO for five consecutive days three weeks after virus injection (**Fig. 1I**), we induced the activation of cortical neurons. The ipsilateral side, with enhanced neuronal activity exhibited an increased number of cFos^+^ neurons (**Suppl. Fig. 4B, C**) forming more contacts with OPC processes compared to the contralateral side of both male and female mice (**Fig. 1J, K**). These results strongly supported the notion that OPCs contact neuronal somata, favouring active neurons, without any sex bias.

To understand whether the augmented OPC-neuron contact was attributed to a greater number of OPCs or branching of OPCs, we performed a morphological analysis of individual OPCs. By administrating a single-dose of tamoxifen to NG2-CreER^T2^ x CAG-^fl^CTA^fl^-EGFP (NG2^CT2-EGFP^) mice (**Fig. 1L**) and applying the same chemogenetic approach to stimulate neuronal activity, we found that ipsilateral OPCs exhibited more branched morphologies and longer processes (**Fig. 1M-O**), without change in soma size (**Fig. 1P**). We also observed increases in OPC density and proliferation (**Suppl. Fig. 4D-F**). These data demonstrated that increases in neuronal activity could stimulate OPC proliferation and morphological changes, thereby augmenting OPC-neuron contacts.

Together, our results strongly suggested that OPCs engage in direct contact with neuronal somata through their processes, with a clear preference for active neurons by increasing their morphological complexity.

### Small neuronal lysosomes were positioned near the OPC-neuron contact site

To identify the physiological significance of the process-somatic contact, we first examined synaptic identity of these sites. At the contact site, OPCs expressed neither presynaptic (vGlut/vGAT) nor the postsynaptic proteins (PSD95), indicating low possibility (if any) of synaptic communication through the process-somatic contact (**Suppl. Fig. 5A-C**).

Considering the correlation between contact frequency and neuronal activity, we explored whether the process-somatic contacts could be linked to neuronal metabolism, particularly focusing on mitochondria and lysosome function. Higher metabolism in neurons often correlates with increased mitochondrial activity and more frequent lysosome release^17, 18^. To investigate the association between OPC-neuron contact and mitochondria or lysosome function, we performed immunostainings using lysosomal membrane marker lamp1, lysosomal content marker Cathepsin D and mitochondria marker Tomm20 (**Fig. 2A, Suppl. Fig. 5D-F**). Our analysis revealed that Lamp1^+^ or Cathepsin D^+^ lysosomes from the neuronal compartment were positioned near the contact site, while mitochondria did not show a similar spatial association (**Suppl. Fig. 5D-F**). Lysosomes exhibit various sizes, with those ready-to-be-released typically smaller and closer to the plasma membrane^19^. To better characterize the lysosomes near the contact site and to demonstrate whether OPC-neuron contact is related to neuronal lysosome release, we performed Lamp1/NeuN/GFP triple immunostaining in NG2^EYFP^ mice and acquired images with STED super-resolution microscopy (**Fig. 2A**). Quantifying the volumes of individual lysosomes of neurons and their shortest distance to the contact sites (**Fig. 2B**), we observed a strong correlation between lysosome volume and its proximity to the OPC surface, with smaller lysosomes positioned closer to the contact site (**Fig. 2C**). To better compare the distance to the OPC surface of small and large lysosomes, we classified lysosomes based on their volume, ranging from 0.02 to 3.0 µm^3^ with a 0.02 µm^3^ interval (**Fig. 2D**). Previously, the volumes of neuronal lysosomes imaged by structured illumination microscopy (SIM) were classified into 7 subgroups^20^, including 0-0.1, 0.1-0.5, 0.5-1.0, 1.0-2.0, 2.0-3.0, 3.0-4.0, 4.0-5.0 µm^3^. Therefore, we divided lysosomes into small (volume <0.1 µm^3^) and large (volume >0.1 µm^3^) subgroups and identified about 35.0% of lysosomes as small (**Fig. 2D**). These small lysosomes were situated closer to the contact site, compared to the large ones (**Fig. 2E**). Furthermore, we compared the volume of lysosomes close to OPCs with those farther away. The shortest distance to the OPC surface was plotted from 0.5 up to 15.0 µm, with 0.5 µm interval (**Fig. 2F**). Lysosomes in proximity to OPCs (distance <2 µm) exhibited smaller volumes than those located away from the contact site (**Fig. 2G**). These data strongly suggested that small lysosomes were located near the contact site, implying the involvement of process-somatic contact between OPCs and neuronal somata in lysosome release.

**Figure 2.**
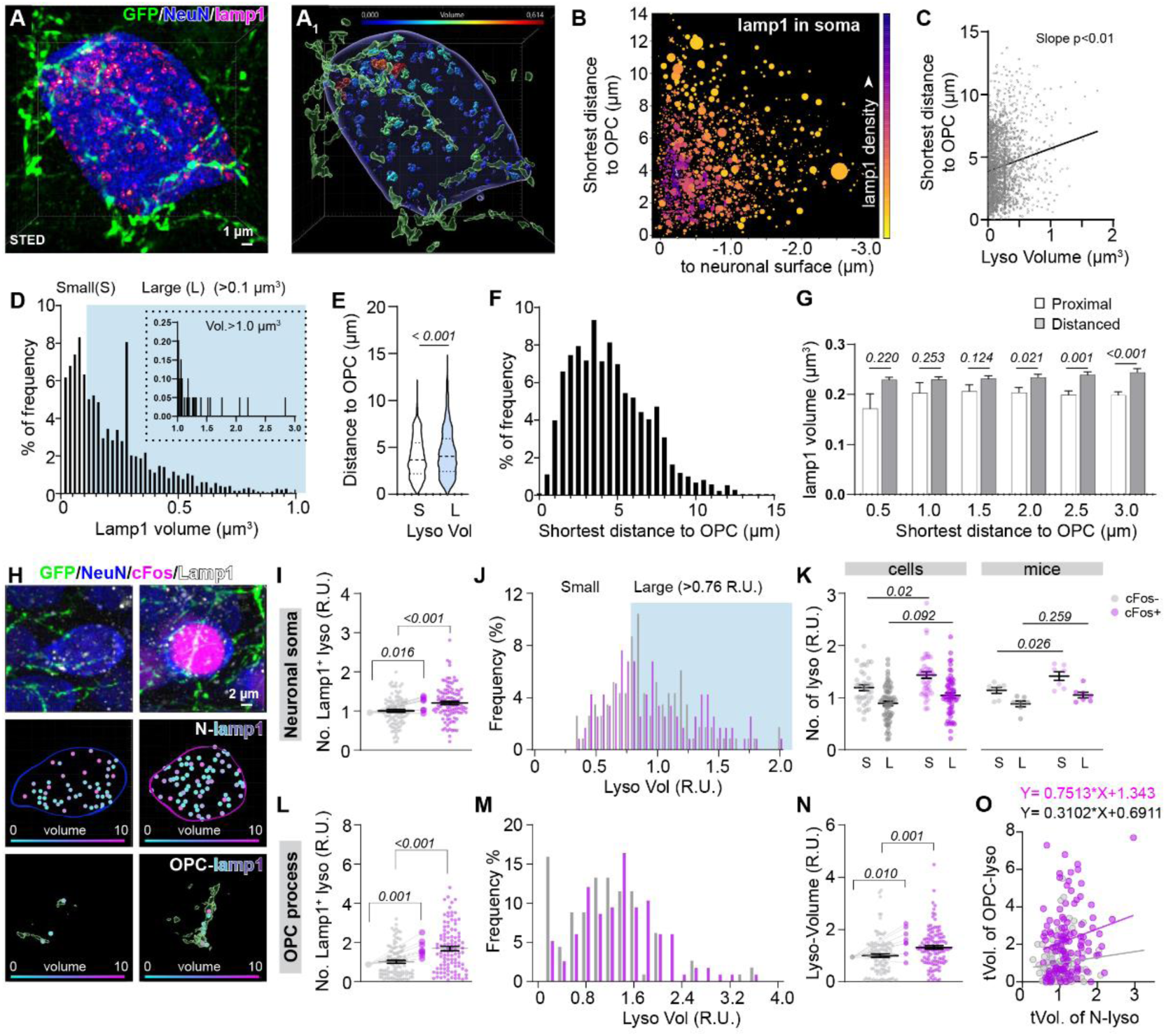
Neuronal lysosomes are positioned close to OPC-neuron contact site. A. Neurons, OPCs and lysosomes were immunolabelled against NeuN, GFP and Lamp1 in the cortex of NG2^EYFP^ mice and 3D reconstructed using Imaris (**A_1_**). NeuN and GFP were imaged with confocal LSM and Lamp1 with a STED microscope. **B** Quantitative analysis of lysosome volume and their shortest distance to the surface of OPCs and neurons (5925 lysosomes from 17 cells of 3 mice). The volume of each lysosome is indicated by the size of the circle and the frequency of the distance between the lysosome and the surface of OPCs and neurons was indicated by the colour code, with the purple indicating higher frequency. **C** Correlation of neuronal lysosome volume and shortest distance to OPC surface (2077 lysosomes from 17 cells of 3 mice). **D** Frequency distribution of lysosome volumes. Lysosomes smaller than 0.1 µm^3^ (35% of total lysosomes) are considered as small. **E** Quantification of the shortest distance between the OPC surface and small or large lysosomes. **F** Histogram of distance between lysosomes and OPC surface. **G** Comparison of the volume of proximal and distal lysosomes to the OPC surface. Lysosomes were classified as proximal or distal to the OPC surface according to their shortest distance to the OPCs. A series of comparison was performed assuming ‘Proximal’ shorter than 0.5, 1.0, 1.5, 2.0, 2.5 or 3.0 µm. **H** Immunostaining of OPCs, lysosomes and neurons with GFP, lamp1 and NeuN in coronal brain slices of NG2^EYFP^ mice. cFos^-^ (left) and cFos^+^ (right) neurons, OPC processes and lysosomes were reconstructed in 3D using Imaris. The volume of lysosomes, both in neurons (N-lamp1) and OPC processes (OPC-lamp1), is shown in different colours from cyan (small) to magenta (large). **I, L** Quantification of the relative number of lysosomes in neuronal soma (**I**) and OPC processes (**L**) contacting neuronal soma at the cellular and mouse level. Lysosome number is normalized to the average lysosome number of cFos^-^ cells. (neuronal soma: 99 cFos^-^ cells and 105 cFos^+^ cells in N=7 mice; OPC processes: 102 cFos^-^ cells and 104 cFos^+^ cells in N= 8 mice; two-sided unpaired t-tests for cells and two-sided paired t-tests for mice). **J, M** Frequency distribution of the relative volume of lysosomes in neuronal soma (**J**) and OPC processes (**M**) contacting neuronal soma at the cellular and mouse level. Lysosome volume is normalized to the average lysosome volume of cFos^-^ cells. Relative volumes less than 0.76 µm^3^ (34.78% of total lysosomes) are considered small. **K** Comparison of the relative volume of small and large lysosomes of cFos^-^ and cFos^+^ neurons at the cellular and mouse level. **N** Comparison of the relative volume of lysosomes in OPC processes contacting cFos^-^ and cFos^+^ neurons (100 cFos^-^ cells and 116 cFos^+^ cells in N=7 mice). **O** Correlation of total lysosomal volume in OPC processes (tVol of OPC-lyso) and that in neurons (tVol of N-lyso) (100 cFos^-^ cells and 116 cFos^+^ cells in N=7 mice). Scale bar in **A**=1 µm, **H**=2 µm.

Given that active neurons release more lysosomes from dendrites and axons^18^, we hypothesized that these neurons might release more lysosomes from the somata as well and recruit more OPC processes to assist in lysosome release. To test this hypothesis, we compared the number of lysosomes in the somata of cFos^+^ and cFos^-^ neurons. Indeed, cFos^+^ neurons exhibited a higher abundance of Lamp1^+^ lysosomes compared to cFos^-^ ones (**Fig. 2H, I**), particularly showing an increase of approximately 20% in the number of small lysosomes in cFos^+^ neurons (**Fig. 2J, K**). Here, the images were acquired with confocal laser scanning microscope, which, unlike STED microscopy, cannot show precise lysosome volume. Hence, we normalized lysosome volume to the mean volume of lysosomes from cFos^-^ neurons and plotted the frequency of each relative volume. Since small lysosomes from STED images (**Fig. 2A**) accounted for about 35.0% of the total, here we designated 35% of the lysosomes from cFos^-^ neurons as small. This classification resulted in lysosomes with a relative unit (R.U.) of <0.76 (comprising 34.78% of the total) being considered as small (**Fig. 2J**). These data suggest that active neurons, particularly those with increased cFos expression, harbour more lysosomes in their somata, especially the small lysosomes.

Furthermore, we observed that the number of lysosomes in each OPC process in contact with cFos^+^ neurons was higher than that in the processes in contact with cFos^-^ neurons (**Fig. 2L**). However, this time the difference could be mainly attributed to the larger lysosomes in the processes (**Fig. 2M**). Additionally, within each process contacting cFos^+^ somata, the average volume of lysosomes was larger than its counterpart contacting cFos^-^ somata (**Fig. 2N**). Consequently, the total volume of lysosomes in proximal OPC processes displayed a close correlation with the total volume of lysosomes in neuronal soma (**Fig. 2O**). These results suggested that OPCs may actively ingest substances released by neuronal lysosomes and subsequently participate in their degradation within their own lysosomal compartments.

The chemogenetic activation of neurons resulted in an increased number, though not an increased volume, of lysosomes in both neuronal somata and OPC processes (**Suppl. Fig. 4G-I**). Moreover, neuronal lysosomes were much closer to the OPC surface in the ipsilateral side. In this context, the total volume of lysosomes in contacting processes and in neuronal somata showed a stronger correlation in the ipsilateral side compared to the contralateral side (**Suppl. Fig. 4K, L**).

In summary, our data demonstrated that active neurons harbour more lysosomes in their somata and receive more contacts from OPC processes, suggesting a putative lysosome-mediated communication between OPCs and neurons.

### Neuronal lysosomes were recruited to OPC-contact sites and engulfed by OPCs

To unravel the physiological significance of the OPC-neuron contact for neuronal lysosomes, we first analysed lysosome trafficking with or without (w/o) OPCs in co-culture by live imaging. Primary neurons, cultured for 9-11 days and visualized by hSyn-hM3D(Gq)-mCherry expression (**Fig. 3A**), were co-cultured with freshly isolated OPCs labelled with Alexa 647 Cell-tracer for an additional two days to allow cellular interaction (**Fig. 3A**). Lysotracker, which labels low pH organelles^21^, was introduced before imaging to visualize lysosomes (**Fig. 3A, B; Suppl. Fig. 6A**). After two days of co-culture, OPCs had already established process-soma junctions with neurons (**Fig. 3B**). Since lysosome release involves trafficking towards the plasma membrane and fusion, we first analysed whether lysosome motility was affected by the presence of OPCs in the culture. In the presence of OPCs, neuronal lysosomes exhibited slower but more efficient movement compared to those in pure neuronal culture (**Fig. 3C**). This was evident from the shorter displacement of each lysosome within 10 minutes and the higher persistence (the rate of displacement/total distance) observed in the co-culture system (**Fig. 3D, E**). This observation suggested a potential regulatory role of OPCs in neuronal activity through the establishment of process-soma contacts, as lysosome trafficking tends to be faster in active neurons^22^. Furthermore, the average distance between each lysosome and its neighbouring lysosomes (to 1, 3, 5 or 9 neighbouring lysosomes) was slightly but significantly decreased within 10 minutes in the presence of OPCs (**Fig. 3F**). These results suggested that lysosomes tend to perform more directed trafficking in the presence of OPCs in the culture.

**Figure 3.**
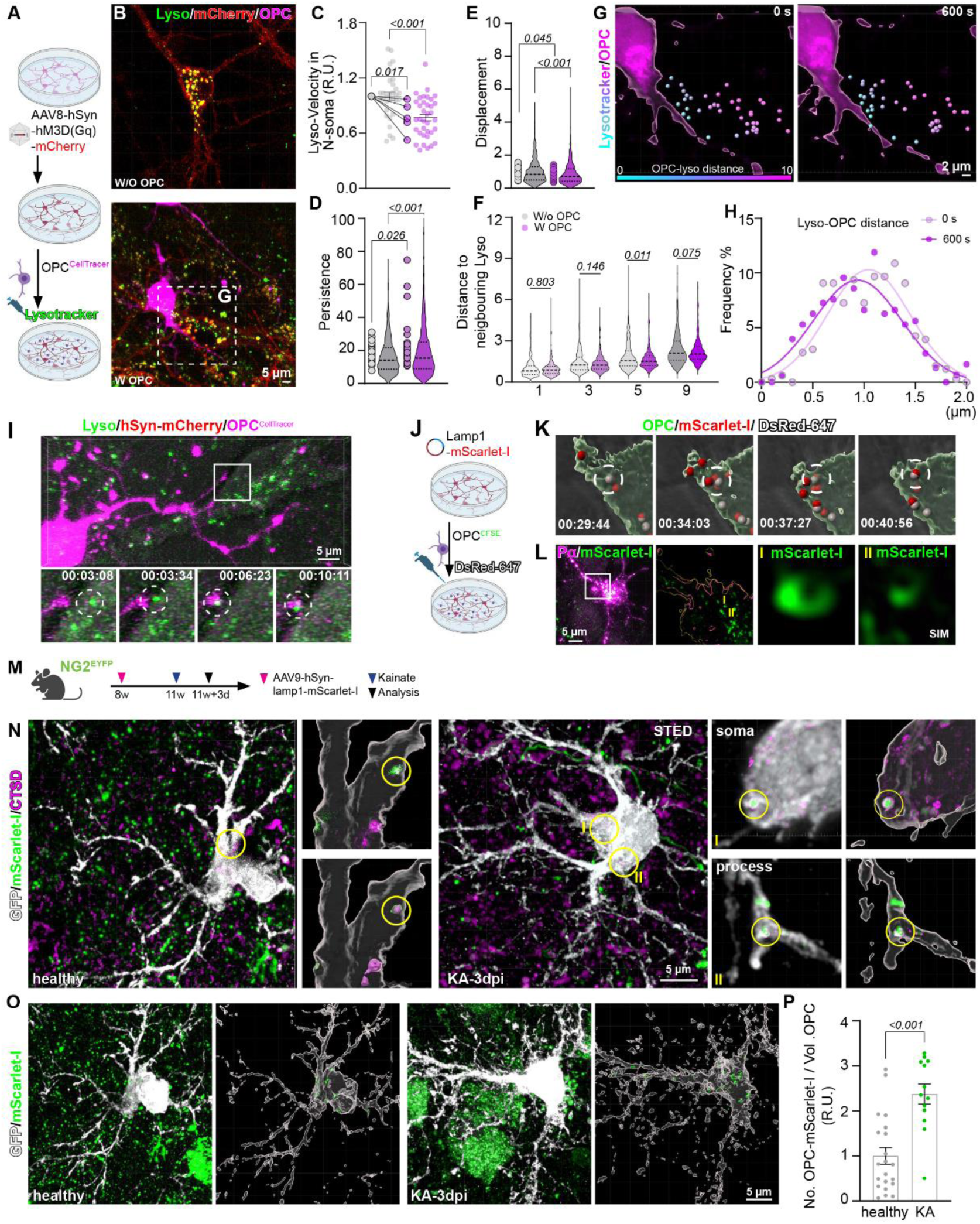
**Neuronal lysosomes are recruited to OPC contact sites and engulfed by OPCs**. **A** Scheme of *in vitro* experiments. **B-H**. **B** Exemplary images of neurons (mCherry^+^) cultured without (w/o) or with (w) OPCs (Alexa647-CellTracer^+^). Lysosomes were visualized by lysotracker. **C-E** Quantification of velocity, movement persistence and displacement of neuronal lysosomes without or with OPCs in culture (**C**: w/o=36 cells and w=31 cells from N=5 independent experiments; **D:** w/o=1253 lysosomes from N=34 cells and w=655 lysosomes from N=23 cells; **E:** w/o=1248 lysosomes from N=34 cells and w=781 lysosomes from N=18 cells). **F** Quantification of the average distance of each lysosome to neighbouring lysosomes in the presence or absence of OPCs within 10 minutes of recording (1: w/o OPCs n=307, w OPCs n=516; 3: w/o OPCs n=307, w OPCs n=516; 5: w/o OPCs n=307, w OPCs n=514; 9: w/o OPCs n=305, w OPCs n=499; two-sided unpaired t-tests). **G** Three-dimensional reconstruction of exemplary images of OPCs and neuronal lysosomes taken at 10-minute intervals. The distance between each lysosome and the OPC surface is indicated by a different colour code, ranging from close (cyan) to far (magenta). **H** Quantification of the distance between the lysosome and OPCs at time 0 (light magenta) and 10 minutes (magenta). **I** Examples of live tracking of the lysosome (green) from the neuronal soma to the OPC processes (magenta). **J** Experimental scheme for **K, L**. **K** Exemplary images showing mScarlet-I-expressing lysosomes co-labelled with Alexa-647 anti-DsRred antibody in OPC processes (green). **L** Cells were post-fixed and the lysosomes were imaged with structured illumination microscope (SIM). **M** Experimental scheme of **N-P**. **N** Brain slices from healthy (left) and the kainate (KA) injected (right) mice were immunostained with GFP (OPC) and Cathepsin D (CTSD, lyososome). mScarlet-I is initially expressed by neuronal lysosomes. GFP and mScarlet were imaged with confocal laser scanning microscope and Cathepsin D (CTSD) was imaged with STED microscope. Three-dimensional reconstruction of the image shows mScarlet-I-expressing lysosomes in OPC processes. **O** Brain slices from healthy (left) and the kainate (KA) injected (right) mice were immunostained with GFP (OPC) and scanned with confocal LSM. **P** Quantification of mScarlet-I^+^ lysosomes in OPC processes in healthy and KA injected cortex (healthy: n=21 cells from N=3 mice; KA: n=13 cells from N=4 mice; two-sided unpaired t-test). Scale bar in **B, I, L, N, P**=5 µm, in **G**=2 µm.

To further elucidate whether the directed trafficking was specifically towards the OPC contact site, we examined the change in the shortest distance between each lysosome and OPC surface within 10 minutes. The results showed a reduction in the distance of individual lysosomes to OPC surface within this time frame (**Fig. 3G, H**), suggesting OPCs likely attract lysosomes and facilitate their release. Indeed, we observed that lysosomes within the neuronal compartment moved towards OPC processes and eventually contacted them (**Fig. 3I**). This result suggested that lysosomes, even in their complete form with low pH, were recruited to the OPC-contact site and potentially taken-up by OPCs. To validate this hypothesis, we transfected neurons with the Lamp1-mScarlet-I plasmid at 5 DIV to visualize neuronal lysosomes. Subsequently, OPCs were added at 10-12 DIV (**Fig. 3J**). To identify whether lysosomes were released as complete organelles or directly engulfed by OPCs before exocytosis, we added anti-DsRed1 antibody conjugated with Alexa-647 to the medium and followed mScarlet-I expressing lysosomes. Since mScarlet-I is expressed on the surface of the lysosome, facing the cytoplasmic side, classical exocytosis leading to the fusion of Lamp1-mScarlet-I to the plasma membrane would not allow the binding of DsRed1 antibody to mScarlet-I. In contrast, in the scenario of a complete lysosomal organelle release, the lysosomal outer membrane would be exposed to the extracellular space, enabling mScarlet-I to be targeted by the Alexa-647-DsRed1 antibody. After about 30 minutes of recording, we observed lysosomes that were positive for both mScarlet-I and Alexa-647, as well as lysosomes solely positive for mScarlet-I, appearing in OPC processes (**Fig. 3K; Suppl. Fig. 6B, C**). These results suggested that OPCs might engulf neuronal lysosomes that were released or located near the plasma membrane. Super-resolution imaging of post-fixed cultured cells by SIM verified the presence of intact neuronal lysosomes in OPC processes, as the mScarlet-I signal (indicating neuronal lysosomes) in OPCs presented a circular pattern (**Fig. 3L**). Taken together, all these data suggested that OPCs could recruit neuronal lysosomes to the contact site by forming physical contact with neurons, and possess the ability to engulf intact neuronal lysosomes.

To further verify the engulfment of neuronal lysosomes by OPCs *in vivo*, we intra-cortically injected AAV9-hSynapsin1-Lamp1-mScarlet-I into NG2^EYFP^ mice and analyzed the mScarlet-I-expressing lysosomes in OPCs three weeks post-injection (**Fig. 3M**). Under physiological condition, a few mScarlet-I-expressing lysosomes were detected in OPC processes (**Fig. 3N-P; Suppl. Fig. 6E**), indicating that the engulfment of neuronal lysosomes by OPC processes occurred with a relatively low probability. However, when neuronal activity was stimulated by cortical injection of kainate (KA), a significantly higher number of lysosomes expressing mScarlet-I were detected in OPC processes and somata three days after KA injection (**Fig. 3N-P; Suppl. Fig. 6E**). The results of the *in vivo* study, consistent with the *in vitro* findings, suggested that the neuronal lysosomes could be engulfed by OPCs through process-somatic contact between OPCs and neurons.

Together, our results demonstrated that OPC processes could form physical contacts with neuronal somata to recruit neuronal lysosomes for exocytosis and eventually engulf them.

### Reduced OPC-neuron contact induces accumulation of neuronal lysosomes and neural circuit change

To elucidate the physiological significance of OPC-mediated neuronal lysosome release and engulfment, we investigated neuronal function in a transgenic mouse model where the branching of OPC processes is reduced (mice with OPC-specific deletion of L-type voltage-gated calcium channels (VGCC) Cav1.2 and Cav1.3)^23^ (**Suppl. Fig. 7A-F**), assuming neurons receive less OPC contacts on their somata in the mutant mice. We induced the double knockout (dKO) of both VGCC genes in OPCs by administering tamoxifen at postnatal day 7 and 8, and analysed the mice at 9 weeks of age (**Fig. 4A**). In the cortex of mutant mice, neurons received less OPC contacts (**Suppl. Fig. 7G, H**), yet the number and the size of their lysosomes were increased compared to control animals (**Suppl. Fig. 7I, J**). Since OPCs preferentially contact more active neurons, we further examined the OPC-contact on cFos^+^ neurons in the mutant mice. As expected, OPC-contacts on cFos^+^ neurons were reduced by about 50% (**Fig. 4B, C**), while the number and the volume of lysosomes in cFos^+^ neurons were increased by about 50% in the mutant mouse brain (**Fig. 4D, E**), particularly with a significant increase in the proportion of large lysosomes (**Fig. 4F, G**). Also, in contrast to the control condition, where neuronal lysosomes were enriched at the contact site, in the mutant mouse brain, lysosome distribution in neuronal somata was scattered with an increased distance to the surface of OPCs (**Fig. 4H, I; Suppl. Fig. 7K**). These results suggested that fewer OPC contacts on neuronal somata led to impairment in lysosome trafficking and aberrant enlargement in lysosomal size. In addition, the ratio between the total volume of lysosomes in neuron-contacting OPC processes versus that in neurons significantly declined in the dKO mouse cortex (**Fig. 4J**), indicating a putative decrease in the engulfing capability of OPCs in the mutant mouse brain. Since lysosomal size negatively correlates with their exocytosis^24^, these results suggested a likely reduction in the exocytosis of neuronal lysosomes in the mutant mouse brain.

**Figure 4.**
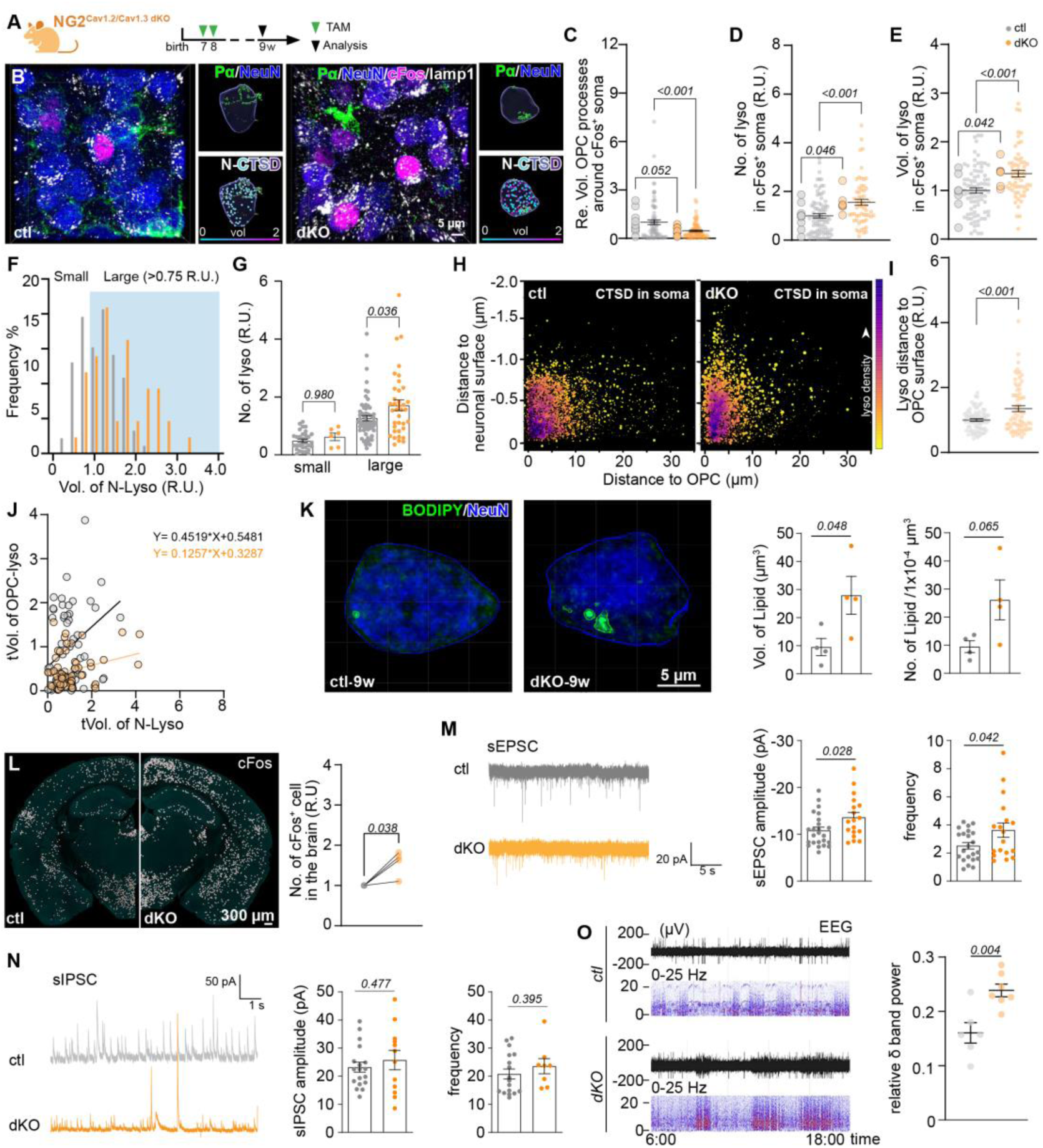
Reduced OPC-neuron contact induces accumulation of neuronal lysosomes and neural circuit change. A. Experimental Scheme. **B** Immunostaining and 3D-reconstruction of OPCs, neurons and lysosomes with PDGFRα (Pα, green), NeuN (blue) and Cathepsin D (CTSD, white) antibodies in control (ctl) and OPC-specific Cav1.2/Cav1.3 double knockout (dKO) mouse brains. Neuronal activity was indicated with cFos immunoreactivity (magenta). **C** Analysis of the relative volume of OPC processes in contact with cFos^+^ neurons in ctl and dKO mouse cortex (ctl: n=74 cells from N=9 mice, dKO: n=74 cells from N=8 mice, two-sided unpaired t-tests). **D, E** Comparison of the relative number and mean volume of lysosomes in cFos^+^ neurons between ctl and dKO mice (ctl: n=84 cells and N=9 mice, dKO: n=59 cells and N=6 mice, two-sided unpaired t-tests). **F** Frequency distribution of the relative volume of lysosomes in neuronal soma of ctl and dKO mice. Lysosomal volume is normalized to the average lysosomal volume of ctl neurons. Relative volumes less than 0.8 relative units (R.U.) (34.78% of total lysosomes) are considered small. **G** Comparison of the relative volume of small and large lysosomes in neurons from ctl and dKO mice. **H** Quantitative analysis of lysosome volume and shortest distance to OPC surface and neuronal surface. The relative volume of each lysosome is indicated by the size of the circle and the frequency of the distance between lysosomes and OPC/neuron surface were indicated by the colour code, with purple indicating higher density (ctl: n=59 cells, dKO: n=59 cells). **I** Relative distance between neuronal lysosomes and OPCs in ctl and dKO mouse brains (ctl: n=5870 lysosomes, dKO: n=4557 lysosomes). **J** Correlation between total lysosome volumes in OPC processes (tVol. Of OPC-lyso) and that in neuronal somata (tVol. of N-lyso) of ctl and dKO mice (ctl: n=73 cells from N=9 mice, dKO: n=68 cells from N=7 mice, two-sided unpaired t-tests). **K** Immunostaining and 3D reconstruction of lipid droplets in neurons of ctl (upper panel) and dKO (lower panel) mouse cortex with NeuN and BODIPY, respectively. Quantification of the number and volume of lipid droplets in the neurons of ctl and dKO cortex. **L** Overview of coronal brain slices from ctl and dKO mice immunostained with cFos and quantification of cFos^+^ cells in the whole brain of both groups (ctl: N=4 mice, dKO: N=4 mice, two-sided paired t-tests). **M, N** Electrophysiological recording of spontaneous excitatory and inhibitory postsynaptic current (sEPSC and sIPSC, respectively) in the cortex of ctl and dKO mice (sEPSC: ctl: n=22 cells from N=4 mice, dKO: n=18 cells from N=4 mice, two-sided unpaired t-tests; sIPSC: ctl: n=18 cells from N=4 mice, dKO: n=12 cells from N=4 mice, two-sided unpaired t-tests). **O** Electroencephalogram (EEG) recordings show significant increase of theta oscillation in the dKO mouse brains (N=6 ctl and 7 dKO mice). Scale bar in **B**=5 µm, in **K**=300 µm, in **L**=50 µm.

Lysosomes, as degradative organelles, play a role in digesting diverse biomolecules, including lipid droplets (LDs). To investigate whether the abnormal accumulation of lysosomes corresponds to lysosomal dysfunction, we examined LD density and volume in cortical neurons of both ctl and dKO mice. This assessment was conducted by employing LD staining combined with NeuN (**Fig. 4K**). The LD volume in neurons of dKO mice was about three times greater than that in the control group, indicating an accumulation of LDs and impaired lysosomal function in dKO mouse neurons. Given that excessive LD accumulation can impact cellular function, we proceeded to analyse neuronal activity and circuits using cFos immunostaining and electrophysiological recordings. In the mutant mouse brain, including cortex, hippocampus, and hypothalamus, the total number of cFos^+^ cells was higher compared to the control group (**Fig. 4L**), implying higher activity of neurons in the mutant mouse brain. Since both excitatory and inhibitory neurons received less OPC contacts (**Suppl. Fig. 8A-C**), we expected a similar change in activity for those neurons. However, we detected elevated mRNA levels of vGlut1 and vGlut2, but not vGAT, in the mutant mouse cortex (**Suppl. Fig. 8D**). In addition, more vGlut1^+^ puncta, but not vGAT^+^ puncta, were observed on neuronal somata, suggesting potentially higher glutamatergic input to the cortical neurons in mutant mouse brains (**Suppl. Fig. 8E**). Electrophysiological recordings substantiated the enhanced local excitatory circuits, as the frequency and amplitude of spontaneous excitatory postsynaptic current (sEPSC) in the mutant mouse cortex were larger (**Fig. 4M**), whereas the inhibitory signals remained unchanged (**Fig. 4N**). This could be attributed to the fact that the majority (94%) of the cFos^+^ neurons are excitatory in the control cortex (**Suppl. Fig. 8G**), making excitatory neurons more affected in the mutant mouse brain compared to inhibitory ones. Nevertheless, as a result, a higher contribution of delta band power (1-4 Hz) to the total brain oscillatory pattern was present in the mutant mouse brain (**Fig. 4O**). These data suggested that OPCs could modulate both neuronal activity and circuits, as well as lipid metabolism, by facilitating neuronal lysosome release and engulfment through process-somatic contact.

### Impaired OPC-mediated neuronal lysosome release was associated with neuronal senescence and neurodegeneration

Increasing evidence links lysosomal dysfunction to neurodegenerative diseases, including, but not exclusively, Alzheimer’s disease (AD)^25^. A recent study using single cell RNA sequencing in early stage AD patients’ brains indicated decreased Cav1.3 (*Cacna1d*) expression in OPCs^26^. Based on this, we hypothesized potential neurodegeneration in Cav1.2/Cav1.3 dKO mice. To explore this possibility, we isolated cortical OPCs from the control and dKO mice at the age of 9 weeks and performed transcriptomic analysis. Genes associated with AD and neurodegenerative diseases were upregulated in dKO OPCs (**Fig. 5A-C; Suppl. Fig. 9A, B**). For example, *Apoe*, one of the major AD risk genes^27^, as well as *Rasgef1a* and *Cd74*, genes indicated to be enhanced in OPCs of AD patients or mouse models, respectively^26, 28^, were also upregulated in dKO OPCs. These observations suggest a connection between Cav1.2/Cav1.3 of OPCs and OPC-mediated neuronal lysosome release/engulfment with neurodegeneration. To validate this hypothesis, we assessed whether neuronal senescence in the adult and aged mutant mouse brain by performing immunostaining of p16INK4A, an established senescence marker^29^ (**Fig. 5D**). Neuronal senescence was similarly low at the age of 9 weeks (data not shown) but increased about 20-fold in the mutant mouse brain during aging (**Fig. 5E**), compared to age-matched control animals. These results suggested that process-soma contact-mediated lysosome release and engulfment could play a protective role against neuronal senescence and degeneration.

**Figure 5.**
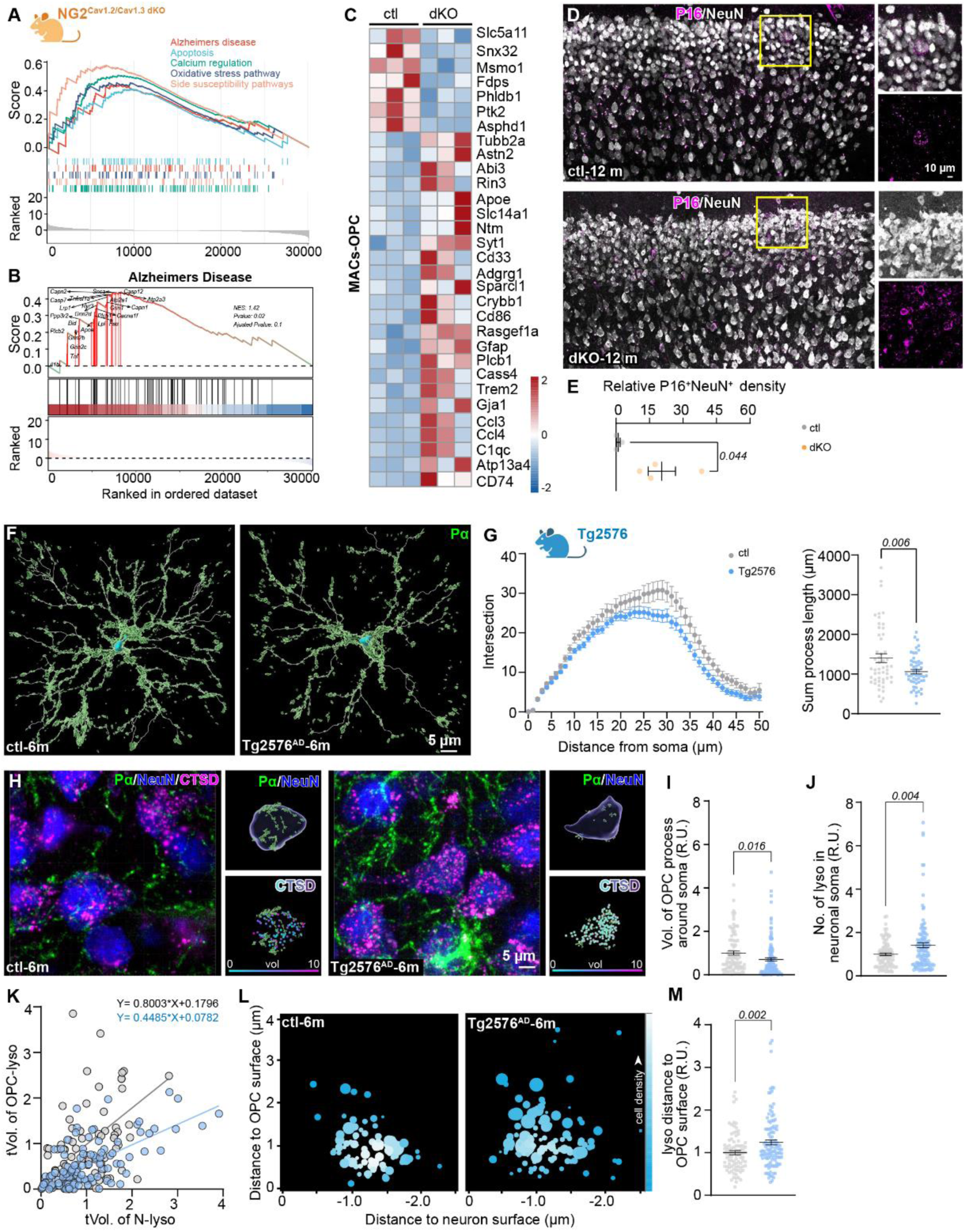
**Impaired OPC-mediated neuronal lysosome release was associated with neurodegeneration**. **A-C** Bulk RNA sequencing of MAC sorted OPCs from control (ctl) and Cav1.2/1.3 double knockout (dKO) cortex show increase in the genes involved in Alzheimer’s disease. **D, E** Immunostaining and quantification of senescent neurons with NeuN (white) and p16INK4A (P16, magenta) in the cortex of ctl and dKO mice at the age of 12 month (ctl: N=3 mice, dKO: N=4 mice, two-sided unpaired t-tests). **F** Three dimensional reconstruction of OPCs immunolabelled with PDGFRα in the ctl and Tg2576 Alzheimer’s disease mouse model at the age of 6 month. **G** Morphological analysis of OPCs from ctl and Tg2576 mice for the number of process intersections and total process length (ctl: n=51 cells, Tg2576: n=52 cells, two-sided unpaired t-tests). **H** Immunostaining and 3D reconstruction of OPCs (PDGFRα, Pα, green), neurons (NeuN, blue) and lysosomes (Cathepsin D, CSTD, magenta) from ctl and Tg2576 mouse cortex. **I** Quantification of the relative volume of OPC processes in contact with neuronal soma in ctl and Tg2576 mice (ctl: n=89 cells, Tg2576: n=116 cells, two-sided unpaired t-tests). **J** Quantitative analysis of lysosome number in neuronal somata in ctl and Tg2576 mouse cortex (ctl: n=87 cells, Tg2576: n=115 cells, two-sided unpaired t-tests). **K** Correlation between total lysosome volume in OPC processes (tVol. of OPC-lyso) and that in neurons (tVol. of N-lyso) of ctl and Tg2576 mice (ctl: n=84 cells, Tg2576: n=114 cells). **L, M** Quantification of the relative mean distance between neuronal lysosomes and the OPC surface in ctl and Tg2576 mice (ctl: n=91 cells, Tg2576: n=117 cells, two-sided unpaired t-tests).

To further substantiate whether OPC-associated neuronal lysosome release is involved in neurodegenerative diseases, we analysed OPC-neuron contact and neuronal lysosomes in an early-stage AD mouse model utilizing Tg2576 transgenic mice. We first analysed OPC morphology in Tg2576 mice at the age of 6 months, representing an early stage of AD pathology^30^, with PDGFRα immunostaining (**Fig. 5F**). OPCs exhibited fewer branches and the total length of processes was shorter in Tg2576 mice (**Fig. 5G**). OPC morphological change was also evident in APP/PS1 AD mouse model (**Suppl. Fig. 9C-E**). We then further analysed OPC-neuron contact and neuronal lysosomes by performing PDGFRα, NeuN and Cathepsin D triple immunostaining (**Fig. 5H**). In the cortex of Tg2576 mice, OPC-contact on neuronal somata was decreased (**Fig. 5I**), despite an increase of lysosome number in neurons (**Fig. 5J**). Consequently, the ratio of total lysosome volume in OPC processes versus that in neurons declined significantly in the cortex of Tg2576 mice (**Fig. 5K**), indicating reduced engulfing and ingesting capability of OPC processes in the AD mouse model. In addition, the average distance between lysosomes and OPC surface was increased in the AD model (**Fig. 5L, M**), indicative of an impaired lysosome trafficking towards the plasma membrane in the cortex of AD mouse model.

Together, these results suggested that impaired OPC-mediated lysosome release and engulfment could be involved in the development of neurodegeneration.

In conclusion, our study highlights the crucial role of OPCs in modulating neuronal lysosome release through direct contact with neuronal somata. The disruption of the process-soma contacts between OPCs and neurons leads to aberrant accumulation of lysosomes and lipid droplets in neuronal soma, disturbance in neural circuits, neuronal senescence, and ultimately neurodegeneration. These findings underscore the significance of OPC-neuron interactions in maintaining proper neuronal function and provide insights into potential mechanisms underlying neurodegenerative conditions.

## Discussion

Lysosomes, responsible for biomolecule degradation and removal of non-recyclable cellular compounds, including misfolded proteins, are typically involved in exocytosis to transport cellular ‘waste’ to the periphery^31^. However, the mechanisms of clearing this ‘waste’ from the extracellular milieu remain unclear. Here, we observed that OPCs form junctions with neuronal somata, where lysosomes, particularly the small and likely ready-to-be-released lysosomes, are located. The number of these contacts was greater in neurons with higher activity. In addition, in these proximal OPC processes, an increased presence of Lamp1^+^ and Cathepsin D^+^ lysosomes were observed, suggesting a lysosome-based signal exchange between neurons and OPCs. Traditionally, lysosome exocytosis involves lysosome trafficking towards and fusion to the plasma membrane, and release of luminal ‘waste’ to the extracellular milieu^32^. In our study, we observed a limited number of neuronal lysosomes in an intact state within OPCs, both *in vitro* and *in vivo* in the mouse brain. This occurrence was more probable when neuronal activity was stimulated. This finding suggests that OPCs might phagocytose lysosomes that are primed for exocytosis or capture lysosomes released into the extracellular space in an intact form. Notably, some neuronal lysosomes, for instance, those identified in cultured OPC processes, were marked with mScarlet (DsRed) antibodies introduced into the culture medium. This indicates a potential, albeit low likelihood that the lysosome surface was exposed to the extracellular environment, a scenario atypical for conventional exocytosis. As OPCs have phagocytic abilities^12, 33^, we cannot rule out the possibility that OPCs unspecifically uptake Alexa647-DsRed-antibody. Through endocytic vesicle trafficking, mScarlet might further fuse with the phagocytosed Alexa647-DsRed-antibody. However, our time-lapse data demonstrates the possibility of direct lysosomal transfer from neurons to OPCs. Further investigations are required to demonstrate how exactly the intact neuronal lysosomes appear in OPC processes.

Recent studies have revealed the capability of OPCs to phagocytose axons and engulf excitatory presynaptic terminals during development^12, 33^. Hence, it is plausible that OPCs might employ similar mechanism to phagocytose membrane-fused neuronal lysosomes by forming physical contacts at the somatic fusion-site. This complex but fine coordination may serve as a protective mechanism for neurons against stress or insults, leveraging the support of glial cells, particularly those renewable ones. When neurons are active, they increase their metabolic rate^34^, resulting in the generation of more ‘waste’ products that need to be transported out of the cell. Failure to effectively export and clear this waste could subject neurons to stress, ferroptosis or degeneration, as seen in lysosomal storage diseases^35–37^. Therefore, having a helper cell capable of cleaning up this ‘waste’ while remaining sustainably available in the brain becomes essential. OPCs emerge as suitable candidates for this role, being the most proliferative cells in the adult and aging brain under physiological condition^38^. Given that not all newly generated OPCs become oligodendrocytes^39^, why should OPCs maintain their proliferative capacity? OPCs are ubiquitously and evenly distributed throughout the brain, each maintaining unique territories^1^. Any kind of OPC loss triggers a fast replenishment via fast proliferation and migration of OPCs^1^. Hence, it is conceivable that OPCs may undergo cell death after taking up neuronal ‘waste’, with this spatial gap filled by newly generated OPCs. Following chemogenetic activation of neurons, the number and total volume of neuronal lysosomes were increased. Subsequently, the ratio of total lysosome volume in OPC processes contacting neurons versus that in neuronal somata was increased, together with OPC proliferation. In contrast, in Cav1.2/Cav1.3 double knockout mice, where this ratio was decreased, OPC proliferation rate was lower compared to the control group^23^. Similarly, in the 2-month-old APP/PS1 mouse model, considered as early-stage AD, OPC proliferation was decreased^40^. Hence, it is plausible that increases in lysosome number and volume in active neurons may lead to an overload of ‘waste’ in OPCs, potentially contributing to OPC death. In normal physiological conditions, the lost OPCs are promptly replaced by newly generated OPCs through the process of proliferation and migration. However, in the context of neurodegeneration, this replenishment mechanism might be impaired, resulting in a deficit in lysosome release and ultimately leading to neuronal senescence. In this study, however, we did not observe any changes in OPC apoptosis in the chemogenetic model. Further investigation is required to address alternative mechanisms of OPC death in this particular scenario. Nevertheless, the delicate balance between neuronal activity, lysosome dynamics, and OPC function appears to be critical for maintaining a healthy and functional neural environment.

In this study, we observed a close correlation between the lysosome volume in OPC processes that contact neurons and the total volume of lysosomes in those neurons. This suggests that OPCs may ingest substances released by neuronal lysosomes, further facilitating degradation within their own lysosomes. Lysosomes play a crucial role in digesting various biomolecules, including lipid droplets that can be broken down into lipid acid and released into the periphery. Lysosomal dysfunction leads to excessive lipid accumulation in cells, impacting cellular function. Impairment in lysosomal function leads to excessive accumulation of lipids in the cells, affecting cellular function. Both lysosomal dysfunction and abnormal lipid accumulation in neurons are intricately connected to neurodegenerative diseases, such as Alzheimer’s and Parkinson’s diseases. For example, before the accumulation of Aβ in the brain of AD mouse model, there is an impairment in the acidification of neuronal autolysosomes^41^. Moreover, both established studies and our own data indicate that OPCs exhibit a reduction in the complexity of their morphology in the brains of early-stage AD mouse models^42^, linking reduced process-somatic communication between OPCs and neurons may contribute AD pathology. Notably, in the Cav1.2/Cav1.3 dKO mutant mice, characterized by reduced arborisation of OPC processes^23^, OPCs exhibited an upregulation of genes associated with neurodegeneration and a significantly larger number of neurons undergoing senescence. These mutant mice also showed enhanced delta power (1-4Hz) in EEG measurements, a phenomenon observed in mild AD mouse model^43^. Additionally, Hmgcs1, a gene involved in cholesterol metabolism, is downregulated in OPCs of early-stage AD patients’ brains^44^ and in Cav1.2/Cav1.3 dKO OPCs. Therefore, OPCs play a crucial role in controlling neuronal lysosomal function and lipid metabolism through direct process-somatic contact. Impairment in this communication at the early stages of neurodegenerative diseases may further drive the detrimental progression of neurodegeneration.

The process-soma contacts between OPCs and neurons may also serve as avenues for signal exchange. Beyond the lysosome’s canonical role in cellular ‘waste’ disposal, more and more studies highlight the role of lysosomes as signalling hubs^45^. Lysosomes contain high levels of ATP, which can be released to the cellular periphery^46^. Coincidently, OPCs express a series of purinergic receptors^47^, including P2Y1^48^, P2X7^48, 49^, P2Y12 receptors, etc. For instance, activation of P2X7 and P2Y1 receptors in OPCs by ATP has been implicated in promoting OPC migration *in vitro*^49, 50^. This raised the possibility that ATP derived from neuronal lysosomes activates P2Y1 and P2X7 receptors, thereby recruiting OPC processes to neuronal somata. However, the temporal sequence of events, whether the process-soma contact precedes lysosome fusion or vice versa, remains unclear. Lysosome tracking in cultured neurons showed that, without OPCs, neuronal lysosomes retain motility and can undergo release. However, in the presence of OPCs in neuronal cultures, lysosome trafficking becomes more directed towards the membrane, suggesting a more efficient lysosome movement facilitated by the OPC contact. Hence, OPC contact emerges as a critical factor for the effective mobilization of lysosomes. Indeed, in the Cav1.2/Cav1.3 double knockout (dKO) mutant mice, where OPC-neuron contact is reduced, aberrant lysosome accumulation in neurons is observed, with lysosomes positioned farther from OPC contact sites.

In conclusion, our study demonstrates that OPCs establish proximal junctions with neuronal somata, which are essential for modulating neuronal lysosome trafficking and activity. This modulation is a critical process in safeguarding neurons against degeneration.

## Supporting information

OPC_Lysosome_Fang_SuppInfo

## Acknowledgement

We thank Hanna Pfeiffer-Unckrich and Harsha Seerapu for their excellent technical support and Daniel Schauenburg for excellent animal husbandry for experimental assistance. We are also grateful to Dr. Manuel Buttini (University of Luxemburg, Luxemburg) for providing brain slices of Tg2576 and corresponding control mice; Prof. Jaqueline Trotter and Prof. Eva-Maria Krämer-Albers (University of Mainz, Mainz, Germany) for providing the NG2 antibody and sharing OPC culture protocol, respectively. This work was supported by grants from the Deutsche Forschungsgemeinschaft (BA 8014/1-1 to X.B.; SPP 1757 to F.K.), University of Saarland (HOMFORexzellent2018 and NanoBioMed Young Investigator grant 2021 to X.B.; HOMFORexzellent2020 and NanoBioMed Young Investigator grant 2020 to H.-F.C.; GradUS global2019 and 2020 to L.-P.F.), the BMBF (EraNet-Neuron BrIE to F.K.), the European Commission (H2020-MSCA-ITN EU-GliaPhD to F.K.) and the Romanian UEFISCDI (PCE 227; PN-III-P4-ID-PCE-2020-2477 to F.K.).

## Author Contribution

L.-P.F. and X.B. conceived the project; L.-P.F., Y.S., C.Y., J.N. and X.B. performed immunostainings and L.-P.F., J.N., A.S. and X.B. performed imaging and analyzed data; C.L. performed cell culture and L.-P.F. performed imaging and data analysis; Y.M. performed STED imaging; H.-F. C. performed SIM imaging; N.Z. performed electrophysiological recordings and data analysis; R.Z. provided technical support and guide for image analysis with Imaris; D.G. performed EEG recordings; Q.G. and X.B. performed intracortical virus injection; A.W. and V.W. prepared viruses; F.K. and X.B. supervised the project; X.B. wrote the manuscript with comments of the other authors.

## Materials and Methods

### Ethics statement

The study was carried out at the University of Saarland in strict accordance with recommendations of European and German guidelines for the welfare of experimental animals. Animal experiments were approved by Saarland state’s “Landesamt für Gesundheit und Verbraucherschutz” in Saarbrücken/Germany (animal license numbers: 65/2013, 12/2014, 34/2016, 36/2016, 03/2021, 07/2021, 08/2021, Perfusion-2023, VM2024-03 and 17/2023; SYXK(YU)2022-0018).

### Animals

All mouse lines were maintained in C57BL/6N background and housed at the animal facility of the CIPMM. Mice were kept on a 12 hour (h) light/dark cycle at 20°C and fed a breeding diet (V1125, Sniff) *ad libitum*. To better observe OPC morphology, NG2-EYFP knock-in mice (NG2^EYFP^) as well as TgH (NG2-CreER^T2^) carrying Rosa26-^fl^STOP^fl^-GCaMP3 (NG2^CT2-GCaMP3^) or CAG-^fl^CTA^fl^-EGFP reporter (NG2C^CT2-EGFP^)^51–54^ were used. To conditionally knock out (KO) Cav1.2 and Cav1.3 in oligodendrocyte precursor cells (OPCs), NG2-CreER^T2^ mice were crossbred to Cav1.2^fl/fl^ (flanking exon 14 and 15 of *cacna1c*) and Cav1.3^fl/fl^ mice (flanking exon 2 of *cacna1d*) (NG2^Cav1.2/Cav1.3 dKO^)^23^. Brain slices of Tg2576 Alzheimer’s disease mouse model was kindly provided by Dr. Manuel Buttini (Neuropathology, Luxembourg Centre for Systems Biomedicine, University of Luxembourg, Luxembourg).

### Tamoxifen administration

Tamoxifen (Carbolution, Neunkirchen, Germany) was dissolved in Miglyol^®^812 (Caesar & Lorentz GmbH, Hilden, Germany) to a final concentration of 10 mg/ml and administered intraperitoneally (100 mg/kg body weight) for two consecutive days at postnatal day (P) 7 and 8 or for five consecutive days at the age of 4 weeks^7, 55^. For single OPC labelling, single-dose tamoxifen was given at the age of 5 weeks.

### Animal surgeries

Adult mice at the age between 8-12 weeks were administered with 10 mg/kg of Carprofen one hour before the surgery. Mouse was fixed on a stereotactic apparatus under continuous inhalational isoflurane (5% for induction and 2% for maintenance with mixture of O_2_ and N_2_O), and the eyes were covered by Bepanthen (Bayer). After sterile cleaning and skin incision, for intracortical injection, the skull was then thinned laterally by 1.5 mm and longitudinally by 1.8-2 mm from Bregma using a dental drill.

***AAV injection*** was introduced to the cortex with coordinates AP:-1.92 mm, ML: 1.5 mm, DV: 1 mm relative to the Bregma. AAV8-hSynapsin1-hM3D(Gq)-mCherry (Addgene, #50474, ≥ 2 x 10^12^ vg/ml) or AAV9-hSynapsin1-Lamp1-mScarlet-I (3.06 x10^12^ vg/ml) in a volume of 0.3 µl was injected intra-cortically with a rate of 0.1 μl/min at a depth of 0.6 mm from pia. The syringe was kept in place for 5 min after the injection to avoid liquid reflux.

***Cortical kainate (KA) injection*** was performed unilaterally following the protocol outlined by Bedner et al.^56^. Briefly, after three weeks of AAV9-hSynapsin1-Lamp1-mScarlet-I injection, 70 nl of 20 mM kainic acid (Tocris) in saline solution were injected into the right hemisphere of anesthetized mice, targeting coordinates AP:-1.84 mm, ML: 1.5 mm, DV: 1 mm relative to the Bregma.

Following the surgeries, the incision was sutured and analgesia was administered intraperitoneally daily for three consecutive days for postoperative pain management. Animals injected with virus or KA also received Tramal (0.4 mg/ml) in the drinking water for the first seven days or Buprenorphin (1 mg/kg) *ad libitum* in the drinking water for the first three days after the injection, respectively.

After three weeks of successful microinjection of AAV8-hSynapsin1-hM3D(Gq)-mCherry, mice were administered with intraperitoneal injection of CNO (5 mg/kg in saline, helloBio) for five consecutive days. Mice were sacrificed 2 hours after the last CNO injection.

***Electroencephalography*** (EEG) was performed using implantable ETA-F10 transmitters (DSI PhysioTel® ETA-F10, Harvard Biosciences, US) as described previously^7^. Briefly, the ETA-F10 transmitter was placed in a subcutaneous pouch in the animal flank and the insulated wires are run subcutaneously to the implantation site on the skull surface (bilaterally 1.6 mm from the sagittal medial axis and 3.4 mm posterior to bregma) and fixed with cyanoacrylate and dental cement (RelyXTM Unicem 2 Clicker, 3M Deutschland GmbH, Neuss, Germany). For precise positioning and drilling, a Stereotaxic Drill Robot (Neurostar, Tübingen, DE) equipped with exchangeable steel round-shaped tip (Ø, 0.9 mm, Hager & Meisinger GmbH, Neuss, DE) was used. The animal skin was closed with an interrupted suture (absorbable suture thread, Eickfil, Eickemeyer, Tuttlingen, DE) and Michel suture clips (F.S.T., Heidelberg, DE). Mice were checked at least once per day and analgesia was administered daily for three consecutive days.

***EEG recording*** was achieved by placing mouse cages on radio-receiving plates (DSI PhysioTel® RPC-1, Harvard Biosciences, US) controlled using the Ponemah Software platform (DSI, St. Paul, US). Video monitoring was performed using the MediaRecorder Software (Noldus Information Technology, Wageningen, NL). Recordings started at least one week after the surgery to enable animal recovery and habituation. Animals were recorded for at least three consecutive days. Data obtained from the telemetric EEG recordings were visualized and analysed using the software NeuroScore (v. 3.3.9318-1, DSI, Data Sciences International, St. Paul, USA). EEG signal was filtered using a 2-200 Hz band pass and a 50 Hz notch filter to remove the power line interference. Absolute spectral powers were calculated by fast Fourier transform using 60 s-long epochs.

### Magnetic activated cell sorting of OPCs

OPCs were sorted with Magnetic activated cell sorting (MACs) technique following the manufacturer’s instructions from Miltenyi Biotec with certain modifications. Mice were perfused with cold Hank’s balanced salt solution lacking Ca^2+^ and Mg^2+^ (HBSS, H6648, Gibco), and cortices were dissected in ice-cold HBSS. After removing debris (130-107-677, Miltenyi Biotec), cells were resuspended in 1 mL of “re-expression medium” containing NeuroBrew-21 (diluted 1:50 in MACS Neuro Medium) (130-093-566 and 130-093-570, Miltenyi Biotec) and 200 mM L-glutamine (diluted 1:100, G7513, Sigma) at 37 °C for 30 min. Subsequently, cells were incubated with Fc-receptor blocker for 10 min at 4 °C (provided with the CD140 MicroBeads kit), followed by a 15 min incubation with a 10 µl microbeads mixture containing antibodies against CD140 (130-101-502), NG2 (130-097-170), and O4 (130-096-670) in a 1:1:1 ratio at 4 °C.

## Primary culture of cortical neurons and OPCs

### Primary cortical neuron culture

Primary neuronal cultures were derived from the cortices of C57BL/6J mice aged P0–1. Cerebral cortices were carefully dissected, and the meninges were removed in ice-cold Earle’s Balanced Salt Solution (EBSS, Gibco). Subsequently, the tissue was finely minced and subjected to digestion with 35 units/mL papain (Worthington, NJ) for 45 min at 37 °C, followed by gentle mechanical trituration. The resulting cell suspension was filtered through a 70 μm cell strainer (Greiner Bio-One). Dissociated cell suspensions were then carefully seeded on 25-mm glass coverslips in 6-well plates (3 × 10^5^ cells/well) or 12-mm glass coverslips in 24-well plates (1 × 10^5^ cells/well). Prior to seeding, the glass coverslips were pre-coated with a solution comprising 17 mM acetic acid, poly-D-lysine (Sigma, St. Louis, MI, USA, P6407), and collagen I (Gibco, A1048301). Neurons were cultured in Neuronal-A (NBA) medium supplemented with 10% FCS, 1% Penicillin/Streptomycin, 1% GlutaMAX, and 2% B-27 supplement (Gibco). On the second day, the medium was replaced with fresh medium to eliminate residual cell debris from the initial cell preparation. Unless specified otherwise, neurons were maintained in the conditioned medium for 8–14 days at 37 °C with 5% CO_2_ before experimental procedures.

### DNA Transfection in Primary cortical neurons

To visualize lysosomes in primary cortical neurons, transfection of neuronal cells was conducted using the DNA calcium phosphate co-precipitation procedure. Prior to transfection, neurons were gently washed twice with NBA medium and fed with serum-free NBA medium without antibiotics. Plasmid DNA was prepared and purified using Endo-free Maxiprep kits (Qiagen, Germany). Subsequently, 4 μg of Lamp1-mScarlet-I (Addgene, #98827) plasmid DNA was combined with 40 μl of 0.25 M CaCl_2_, followed by the dropwise addition of an equal volume of 2x HBS buffer. The resulting calcium phosphate-DNA solution was incubated for 20 min at room temperature (R.T.). This solution was then gently distributed over the neurons at 7 days *in vitro* (DIV) and this amount is suitable for one well of the 6-well plate or three wells of the 24-well plate. After careful swirling, neurons were incubated with the mixture for 30 min at 37 °C with 5% CO_2_. Following the addition of DNA, the culture medium was aspirated, and the cells were rinsed twice with PBS before being replenished with fresh NBA culture medium. mScarlet-I began to be expressed by the neurons approximately 5 days post-transfection.

### OPC sorting for cell culture

OPCs were obtained from neonatal C57BL/6N or NG2^EYFP^ mouse brains (age < P7) using a modified MACs procedure. Briefly, cortices were dissected in cold HBSS solution (without Ca^2+^/Mg^2+^), followed by dissociation using the Neural Tissue Dissociation Kit (130-092-628, Miltenyi Biotec). The dissociation process was stopped by adding 10 ml of DMEM high glucose medium (11965092, Fisher Scientific) with 1% horse serum (HS, Fisher Scientific). After a brief centrifuge, cells were resuspended in DMEM and filtered through 70 µm and 40 µm cell strainers successively. The collected cells were then resuspended in basic medium (NeuroBrew-21 in MACs Neuro Medium (1:50) (130-093-566 and 130-093-570, respectively, Miltenyi Biotec) with 200 mM L-glutamine (1:100, G7513, Sigma)). The cells were incubated at 37°C for 1.5-3 hours (hr), followed by centrifugation at 4°C and resuspension with 134 µl/brain of DMEM+1% HS, along with a mixture of microbeads (20 µl/brain) conjugated with antibodies against CD140 (130-101-502), NG2 (130-097-170), and O4 (130-096-670) in a 1:1:1 ratio for 15 min at 4 °C. The cells were then resuspended in 2 ml DMEM+1% HS/brain, followed by centrifugation at 1200 rpm for 10 min at 4°C. In subsequent steps, DMEM+1% HS was used instead of MACs buffer, following the manufacturer’s instructions (130-101-502). Finally, 5 ml of proliferation medium (containing 4.2 µg/ml Forskolin (F6886, Sigma), 10 ng/ml CNTF (450-13, Pepro Tech), 10 ng/ml PDGF-AA (100-13A, Pepro Tech), 1 ng/ml NT-3 (450-03, Pepro Tech) in basic medium) was added to flush out OPCs. OPCs isolated from C57Bl/6N were further labeled with CellTracer CFSE (C34570, Invitrogen) or CellTracer FarRed (C34554, Invitrogen). Cells were seeded in plates pre-coated with Poly-L-Lysine (Merck) at a density of 1×10^5^ cells/well (24-well plate) or 3×10^5^ cells/well (6-well plate).

### OPC-neuron co-culture

To investigate lysosome communication between OPCs and neurons, a co-culture approach was employed. Neurons and OPCs were plated together under optimized conditions, combining proliferation medium and neuron NBA culture medium at a 1:1 ratio, resulting in 1 ml of medium per 24-well and 2 ml per 6-well plate. At 10-12 DIV of neuronal culture, OPCs were seeded at a density of 5 x 10^4^ cells/well of 24-well plate or 1.5 x 10^5^ cells/well in 6-well plate on top of the neurons. The co-cultures were maintained in optimized medium for additional 2-3 days throughout the experiments.

### Immunohistochemistry

Mice were perfused with PBS followed by 4 % PFA. For NG2 immunostaining, we optimized the fixation protocol by perfusing animals with 2 % paraformaldehyde and post-fixation for four hours at 4°C. After post fixation in 4 °C overnight, coronal slices in 40 µm thickness were prepared using a Leica VT1000S. After 1 hr incubation with blocking buffer (5% HS with 0.5% Triton in PBS), free floating slices were incubated with primary antibodies (Suppl. Table 1) at 4 °C overnight, followed by secondary antibody (Suppl. Table 2) incubation at R.T. for 2 hrs at the next day. DAPI (25 ng/ml) was used for general stain of nuclei (A10010010, Biochimica). For lipid staining, slices are secondary antibody washing, were incubated with BODIPY^TM^ 493/503 (Invitrogen, D3922, 1mg/ml in DMSO) diluted 1:5000 with PBS for one hour. After three washes, the slices were mounted for further analysis.

The brains of APP/PS1 mice and the age-matched control animals at 6-month-old were cryoprotected in 30% sucrose, embedded with Tissue-Tek O.C.T. Compound (Sakura, 4583), and processed for 20 µm cryosections. Sections were permeated in 0.5% Triton-X100 in PBS. Sections were blocked with 2% bovine serum albumin (BSA), and proceeded for primary and secondary antibody incubation as described above.

***Antigen retrieval treatment*** was introduced to enhance the signals of cathepsin D immunostaining in Tg2576 transgenic and control mice^5^. Prior to the blocking step, the following procedures were carried out: Heat treatment in citrate buffer (pH=6.0): Brain slices mounted on the object slides were kept in an appropriately sized container filled with citrate buffer (10 mM citric acid and 0.05% Tween 20). The container was immersed in a water bath at a temperature exceeding 95°C for 30 min. After cooling to R.T., the slices were subsequently processed for the immunostaining procedure.

**Supplementary Table 1.** List of primary antibodies.

| **Antibodies** | **Host** | **Dilutions** | **Cat. No.** | **Company** |
| --- | --- | --- | --- | --- |
| **Cathepsin D** | Rabbit | 1:500 | ab75852 | abcam |
| **cFos** | Guinea pig | 1:1000 | 226004 | Synaptic Systems |
| **CTIP** | Rat | 1:500 | 650601 | Biolegend |
| **GABA** | Rabbit | 1:500 | 20094 | Immunostar |
| **GAD67** | Mouse | 1:500 | MAB5406 | Millipore |
| **GFP** | Goat | 1:1000 | 600-101-215 | Rockland |
| **GFP** | Rabbit | 1:1000 | 632593 | Clontec |
| **IBA1** | Rabbit | 1:1000 | 019-19741 | Wako |
| **IBA1** | Goat | 1:500 | ab5076 | abcam |
| **Kir4.1** | Rabbit | 1:250 | APC-035 | Alomone Labs |
| **Lamp1** | Rat | 1:250 | 121602 | Biolegend |
| **Nav1.2** | Rabbit | 1:250 | ASC-002 | Alomone Labs |
| **NeuN** | Rabbit | 1:500 | ab104225 | abcam |
| **NeuN** | Mouse | 1:250 | MAB377 | Millipore |
| **NG2** | Rat | 1:100 |  | Trotter Lab |
| **p16INK4a** | Mouse | 1:250 | MA5-17142 | Invitrogen |
| **Parvalbumin** | Mouse | 1:500 | P3088 | Sigma |
| **PDGFRα** | Goat | 1:500 | AF1062 | R&D Systems |
| **PSD-95** | Rabbit | 1:250 | 51-6900 | Thermo Fisher |
| **SATB2** | Rabbit | 1:500 | ab92446 | abcam |
| **Somatostatin** | Rat | 1:250 | MAB354 | Millipore |
| **Synaptophysin** | Mouse | 1:250 | S5768 | Sigma |
| **TBR1** | Rabbit | 1:500 | 49661 | Cell Signaling |
| **Tomm20** | Rabbit | 1:250 | ab78547 | abcam |
| **VGAT** | Rabbit | 1:250 | 131002 | Synaptic Systems |
| **VGLUT1** | Guinea pig | 1:250 | 135404 | Synaptic Systems |

**Supplementary Table 2.** List of secondary antibodies.

| Antibodies | Fluorophore | Dilutions | Cat. No. | Company |
| --- | --- | --- | --- | --- |
| Donkey anti-mouse | Alexa Fluor® 488 | 1:1000 | A21202 | Thermo Fisher |
|  | Alexa Fluor® 546 | 1:1000 | A10036 |  |
|  | Alexa Fluor® 647 | 1:1000 | A31571 |  |
|  | DyLight® 755 | 1:1000 | SA5-10171 | Invitrogen |
| Donkey anti-rabbit | Alexa Fluor® 488 | 1:1000 | A21206 | Thermo Fisher |
|  | Alexa Fluor® 546 | 1:1000 | A10040 |  |
|  | Alexa Fluor® 647 | 1:1000 | A31573 |  |
|  | Alexa Fluor® 790 | 1:1000 | A11374 |  |
| Donkey anti- goat | Alexa Fluor® 488 | 1:1000 | A11055 | Thermo Fisher |
|  | Alexa Fluor® 546 | 1:1000 | A11056 |  |
|  | Alexa Fluor® 647 | 1:1000 | A21447 |  |
|  | Alexa Fluor® 750 | 1:1000 | ab175744 | abcam |
| Donkey anti-rat | Alexa Fluor™ Plus 405 | 1:250 | A48268 | Invitrogen |
|  | DyLight-755 | 1:500 | SA5-10031 | Thermo Fisher |
| Donkey anti- guinea pig | Alexa Fluor® 488 | 1:500 | 706-545-148 | Jackson Immuno Research Europe Ltd |
|  | Alexa Fluor® 647 | 1:500 | 706-605-148 |  |
| Goat anti-rat | ATTO® 647N | 1:250 | 40839 | Sigma-Aldrich |
| Goat anti- rabbit | ATTO® 647N | 1:100 | 612-156-120 | Rockland |

## Image acquisition and analysis

Whole brain slices were scanned with fully automated slide scanner AxioScan.Z1 (Zeiss, Jena) at the AxioScan core facility in CIPMM and cell density was analysed manually using ZEN software (Zeiss, Jena).

### Live imaging of lysosomes in vitro

To track lysosome trafficking, lysosomes were labelled with Lysotracker (1:2000, L7526, Invitrogen) introduced to the medium immediately prior to the imaging or with Lamp1-mScarlet-I transfected to neurons as described above. To identify whether lumen side or the surface of lysosome is exposed to the extracellular space or not, either Alexa Fluor® 647 Anti-DsRed (self-conjugated with Alexa Fluor® Antibody Labeling Kits (A20186, Invitrogen) and Living Colors® DsRed Polyclonal Antibody (632496, Takara) or Alexa Fluor® 647 Anti-LAMP1 (ab237307, Abcam) were added to the culture medium at a concentration of 1:1000 immediately prior to live imaging.

Cells were placed in the imaging chamber of LSM 780 confocal microscope with consistent temperature (37 °C) and CO_2_ supply (5%). Live imaging was performed with 63× objective (NA.1.4) for 2-5 µm z-stack with 1 μm interval. Two to four channels were imaged simultaneously with 0.8 Hz, averaging with 2-4 acquisition, for 10-120 min. Images were analysed with Imaris software.

### OPC morphology analysis

Images were acquired with LSM 880 confocal microscope (Zeiss, Oberkochen) with 63x objectives (N.A. 1.4, Oil) with a 1 μm interval. OPC morphology was analyzed with ‘filament function’ of Imaris (Version 9.6, Oxford Instruments) with following settings: ‘autopath’ selection with a maximum diameter of 10 µm and seed points with 0.3 µm; elimination of seed points within a 20 µm sphere region around starting points; and removal of disconnected segments using a 0.6 µm smooth method. Subsequent to filament reconstruction, individual datasets for Sholl analysis were exported into distinct Excel files for further examination.

### Analysis of process-soma junction

Images were acquired with LSM 880 confocal microscope (Zeiss, Jena) with 63x objectives with 1 μm interval. Three D-reconstruction of neuronal soma was conducted with Imaris. Subsequently, OPC processes were reconstructed in ‘surface’ mode, touching objects were splited with 0.5 µm seed point diameter using morphological split, and the distance to neuronal soma was set as <0.25 µm. Following OPC process reconstruction, individual volume of contacting segment of each processes were exported into separate Excel files for further analysis.

### Lysosome analysis

After completing OPC and neuronal somata surface reconstruction, the surface reconstruction was employed as templates for spot reconstruction. The following settings were applied: different spot sizes with an estimated XY diameter of 0.5 µm. Following spot reconstruction, separate datasets for volume and the shortest distance to the OPC and neuronal surfaces were exported into distinct Excel files for detailed analysis.

While for lysosomes imaged with super resolution STED microscope, the absolute volume of the lysosome was quantified, the relative volume of lysosomes was quantified for confocal microscope images with following method: Initially, the average volume of all lysosomes within each cell was assessed. Subsequently, we determined the average volume of lysosomes in cFos^-^ or ctl cells. Finally, we normalized the volume of lysosomes of cFos^+^ cells (of neurons of dKO mice) to the average value of cFos^-^ cells (or neurons of ctl cells).

### Structured illumination microscopy (SIM)

To elucidate the structure of neuronal lysosomes appeared in OPCs, co-cultured Lamp1-mScarlet-I transfected neurons and OPCs were fixed, mounted using the mounting medium (Immu-Mount, epredia) and examined utilizing a SIM microscope (Zeiss Elyra PS.1). Z-stack images in 200 nm z-stack were acquired with 63x objective (1.4 Oil DIC M27), employing three gratings to scan the fine regions of the contacted sites between neuron soma and OPC processes. All acquired images underwent post-processing to achieve a resolution of 100 nm in x and y dimensions and 200 nm in z dimension.

### Stimulated emission depletion (STED) microscopy

For STED microscope imaging, secondary antibodies of ATTO series (**Table 2**) were used. Imaging was performed on an inverted STED Microscope (Expert Line, Abberior Instruments, Göttingen, Germany) using 488 nm, 561 nm, and 640 nm pulsed excitation lasers with respective detections at 498–520 nm, 605–625 nm, and 650–720 nm. Brain slices were visualized with a 100x silicone oil immersion objective (NA 1.4, UPLSAPO100XS, Olympus, Hamburg, Germany). STED images of lysosomes were acquired with 640 nm excitation and 775 nm STED laser with a toroidal (“donut”) depletion pattern of the STED focus. The images were recorded using Imspector software (Abberior) with a voxel size of 20×20×300 nm^3^. The pinhole size was set to 1.08 Airy Units. For 3D rendering with Imaris, image stacks were planewise linearly deconvolved (Wiener filtered) using theoretical point-spread functions and customized MATLAB codes, and further denoised with Gaussian and Median filters.

## Electrophysiology

Mice were anesthetized with isoflurane and decapitated. Brain was swiftly extracted and immersed in ice-cold, oxygenated (with 5 % CO_2_ and 95 % O_2_) solution with a composition of (in mM) 87 NaCl, 3 KCl, 25 NaHCO_3_, 1.25 NaH_2_PO_4_, 3 MgCl_2_, 0.5 CaCl_2_, 75 sucrose, and 25 glucose with pH 7.4. Semi-sagittal brain slices in 300 µm thickness were prepared with vibratome (Leica VT 1200S, Nussloch, Germany). Slices were then carefully transferred to a nylon basket slice holder for subsequent incubation in artificial cerebral spinal fluid (ACSF) at 32°C for 30 minutes. The ACSF composition consisted of (in mM) 126 NaCl, 3 KCl, 25 NaHCO_3_, 15 glucose, 1.2 NaH_2_PO_4_, 2 CaCl_2_, and 2 MgCl_2_. Following this incubation period, the slices were maintained at room temperature with continuous oxygenation until further use.

Semi-sagittal brain slices were carefully transferred to a recording chamber and continuously perfused with oxygenated ACSF containing 1 mM MgCl_2_ and 2.5 mM CaCl_2_ at a flow rate of 2–5 mL/min. To isolate spontaneous excitatory postsynaptic currents (sEPSCs), 50 μM strychnine and 50 μM picrotoxin were added to block inhibitory synaptic transmission. Pyramidal neurons were morphologically identified using an Axioskop 2 FS mot microscope (Zeiss, Jena, Germany) equipped with a 40x water immersion objective and a QuantEM 512SC camera (Photometrics, Tucson, USA). Whole-cell membrane currents were recorded using an EPC 10 USB amplifier (HEKA, Lambrecht, Germany), low pass filtered at 3 kHz, and data acquisition was managed by Patchmaster software (v2×90.5, HEKA). Patch pipettes (7–9 MΩ) were pulled from borosilicate capillaries (outer diameter: 1.5 mm; Sutter, USA) using a Micropipette Puller (Model P-97, Sutter Instrument Co., CA). The pipettes were filled with an internal solution consisting of (in mM) 125 cesium gluconate, 20 tetraethylammonium (TEA), 2 MgCl_2_, 0.5 CaCl_2_, 1 EGTA, 10 HEPES, and 5 Na_2_ATP (pH 7.2). Spontaneous excitatory and inhibitory postsynaptic currents (sEPSCs and sIPSCs) of pyramidal neurons in the medial prefrontal cortex (mPFC) were recorded for 40 s in voltage-clamp mode at holding potentials of –70 mV and +30 mV, respectively. Currents above 10 pA were analyzed using MATLAB.

The data generated by PatchMaster were imported into MATLAB (MathWorks, MA, USA) using a module adapted from sigTOOL. Subsequently, evoked EPSC and IPSC traces from identical cells underwent manual verification and were combined. The averages of EPSC/IPSC traces from each cell were then employed for analysis. Custom MATLAB routines were utilized for data analysis^57^.

## Quantitative real time PCR

The brain tissue were homogenized using the previously outlined procedure^58^. Subsequently, mRNA extraction was carried out utilizing the NucleoSpin RNA Plus XS kit (740990.50, Macherey-Nagel). Reverse transcription was accomplished using the Omniscript kit (205113, QIAGEN). RT-PCR was conducted employing the EvaGreen kit (27490, Axon) with CFX96 Real Time System (BioRad). The primer sequences used for qRT-PCR are provided in **Table 3**.

**Table 3.**
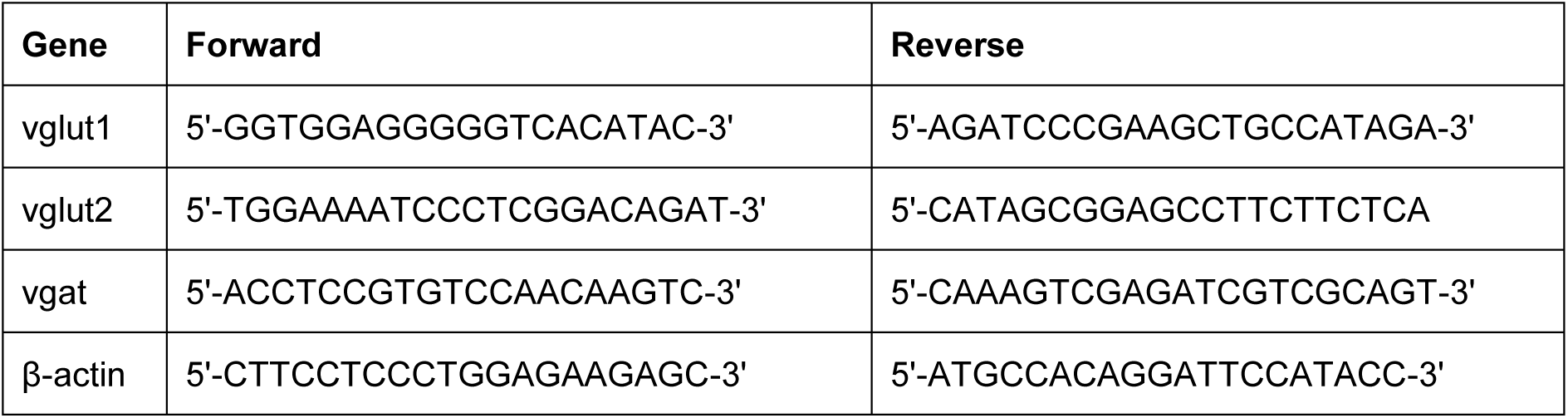
Primers used for qRT-PCR.

## Next-generation RNA sequencing

The library underwent preparation and sequencing procedures conducted by Novogene, employing a series of meticulous methods. Initial quality assessment involved 1% agarose gel electrophoresis to evaluate RNA degradation and potential contamination. Subsequently, sample purity and preliminary quantitation were determined using the Bioanalyser 2100 from Agilent Technologies, USA. This instrument was also instrumental in assessing RNA integrity and final quantitation.

For library preparation, oligo d(T)25 magnetic beads were utilized to selectively isolate mRNA from the total RNA sample, employing a method known as polyA-tailed mRNA enrichment. Following this, mRNA underwent random fragmentation, and cDNA synthesis ensued using random hexamers and the reverse transcriptase enzyme. Upon completion of the first chain synthesis, the second chain was synthesized with the addition of an Illumina buffer. Through the presence of dNTPs, RNase H, and polymerase I from E. coli, the second chain was obtained via nick translation. The resultant products underwent purification, end-repair, A-tailing, and adapter ligation. Fragments of the appropriate size were enriched through PCR, introducing indexed P5 and P7 primers, with final products subjected to purification.

Verification of the library was carried out using Qubit 2.0 and real-time PCR for quantification, while the Agilent 2100 bioanalyzer was employed for size distribution detection. Quantified libraries were pooled and subsequently sequenced on the Illumina Novaseq 6000 platform, based on effective library concentration and data volume.

The qualified libraries underwent Next Generation Sequencing (NGS) utilizing Illumina’s Sequencing Technology by Synthesis (SBS), wherein fluorescence detection was employed for nucleotide identification during the synthesis of the complementary chain. The Novaseq 6000 sequencing system was instrumental in conducting the parallelized and massive sequencing of the libraries. The sequencing strategy employed was paired end 150bp (PE150).

## RNA-seq data processing

The quality of RNA sequencing reads was assessed through FastQC (https://www.bioinformatics.babraham.ac.uk/projects/fastqc/). Alignment to the GRCm38 Mus musculus genome was carried out using HISAT2 v2.0.5^59^ with default parameters. The gene count matrix for each sample was generated using featureCounts v1.5.0-p3^60^. Subsequent analysis utilized the DEseq2 v1.20.0 package^61^ in R program. Genes with a normalized count below 10 were excluded from downstream analysis. Significantly deregulated genes were identified with a false discovery rate below 0.05.

Differential expression analysis was performed with the ‘DESeq’ function with default parameters, and log fold change shrinkage was applied to the analysis results. Heatmaps depicting differentially expressed genes (DEGs) with a p value < 0.02 were visualized using pheatmap v1.0.12^62^. Selected gene set enrichment analysis for Gene Ontology (GO) and Kyoto Encyclopedia of Genes and Genomes (KEGG) pathways was conducted with ClusterProfiler v3.8.1^63^.

## Statistical analysis

Data were analyzed with Graphpad Prism 10.1.2 and Originpro 2022, and figures were generated with Adobe Indesign 2023/2024 and Adobe illustrator 2023/2024. For all immunostainings, two randomly selected brain slices of each mouse were used. At least three animals of each sex were analyzed per group and the data from both sexes were pooled as no sex-difference was observed. The used statistical analysis is indicated in the figure legends. Normal distribution was tested within Graphpad Prism, and normally distributed dataset were analyzed with unpaired t-tests, paired t-test, one-way ANOVA and two-way ANOVA (indicated in each figure legend), while the Kruskal-Wallis test was used for non-normally distributed datasets. P-values are indicated in the figures. In **Fig. 3H**, non-linear curve fitting was employed. A simple linear regression analysis was conducted to assess the relationship between the sum lysosome volume in the respective OPC and neuronal somata. Data were shown as mean ± SEM.

