## Supplementary material for "Neuronal lysosome transfer to oligodendrocyte precursor cells: a novel mechanism of neuron-glia communication and its role in neurodegenerative diseases": OPC_Lysosome_Fang_SuppInfo

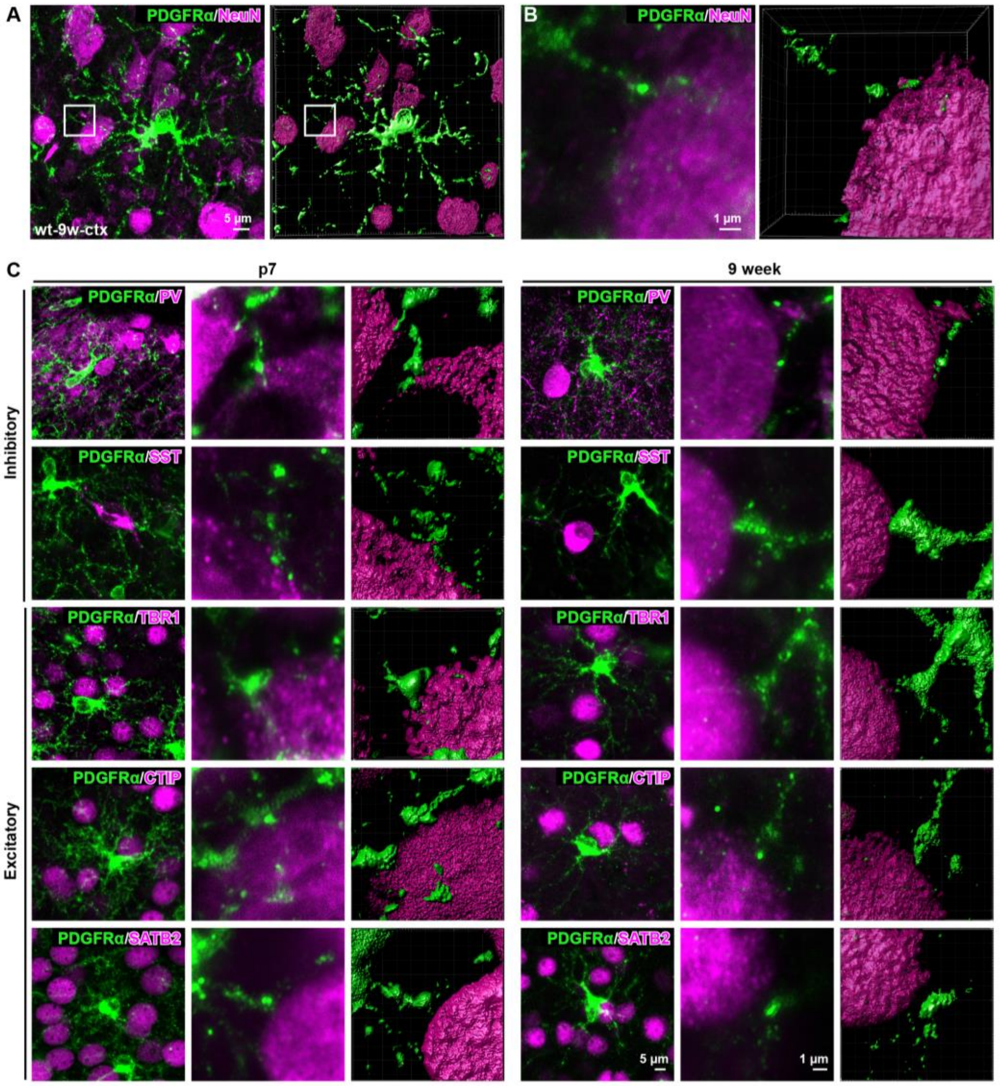


**Supplementary Figure 1. OPCs form proximal junction with neurons in the cortex during development and in the adult**. **A, B** Immunostaining of OPCs and neurons with PDGFRα and NeuN in the cortex of adult (9-week) wildtype mouse. **C** Immunostaining of OPCs and various types of neurons combining PDGFRα with Parvalbumin (PV), somatostatin (SST), TBR1, CTIP, SATB2 in the developing and adult brain. Scale bar in **A** and **B**=5 µm, in **C**=5 or 1 µm, as indicated.

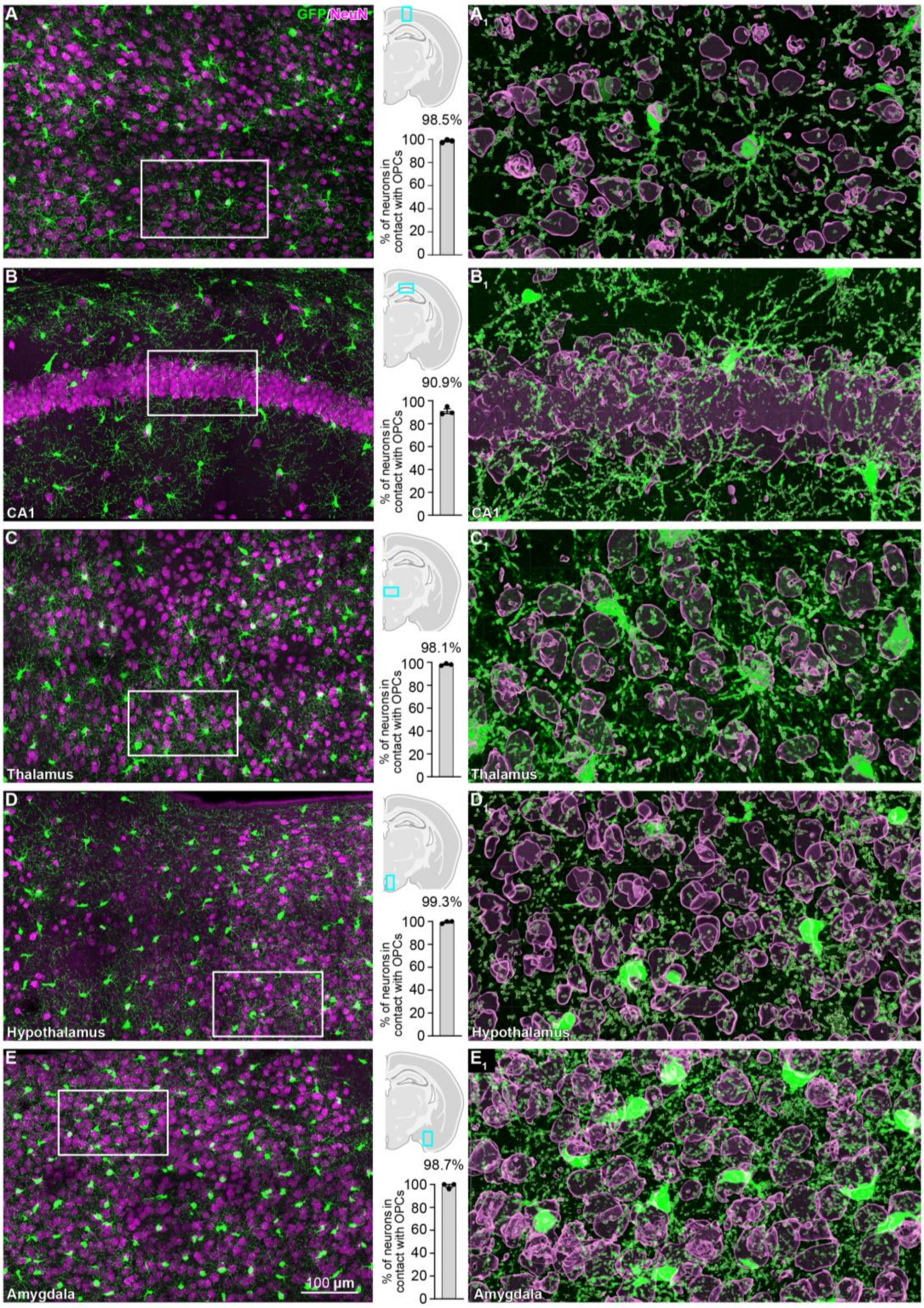


**Supplementary Figure 2. OPC-neuron junction is common feature in all grey matter regions.** Immunostaining of OPC-neuron contact immunostained with GFP and NeuN from cortex (**A**), hippocampal CA1 region (**B**), thalamus (**C**), hypothalamus (**D**) and amygdala (**E**) of NG2$^{EYFP}$ mice. Percentage of neurons receiving OPC contact was quantified.

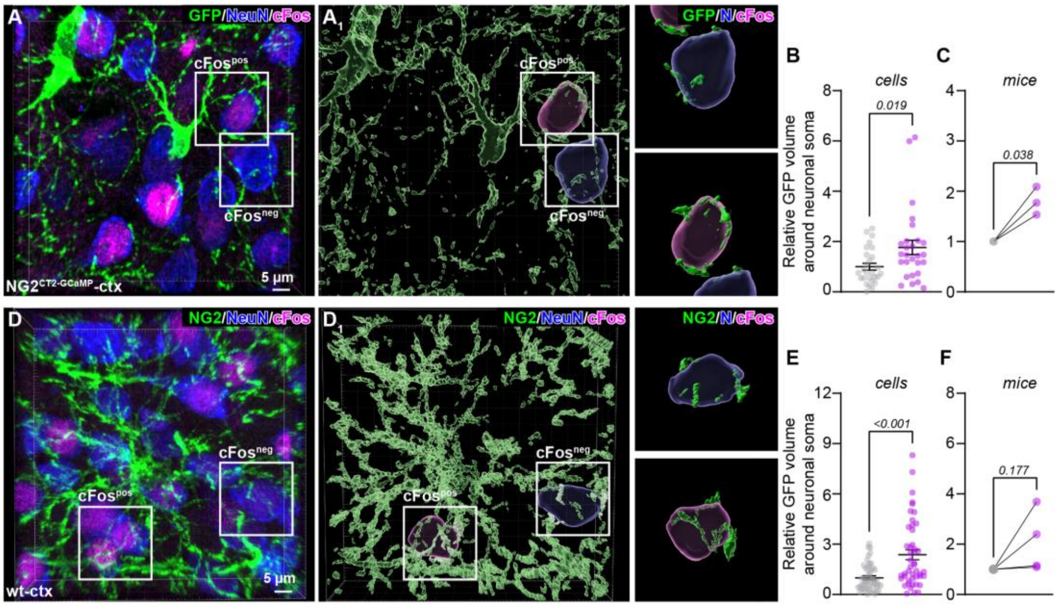


**Supplementary Figure 3. OPCs preferentially form contact with active neurons. A, D** Immunostaining of OPCs with GFP (**A**) or NG2 (**D**) and neurons with NeuN in the cortex of NG2-CreER$^{T2}$ x GCaMP3 (NG2$^{GCaMP3}$) mice or wildtype mice, repectively. Tamoxifen was administered to NG2$^{CT2\text{-}GCaMP3}$ mice at the age of 5 weeks for five consecutive days and analyzed at 9w. **B, C, E, F** Quantification of relative volume of OPC processes (GFP$^{+}$) wrapping cFos$^{+}$ and cFos$^{-}$ neurons. Scale bar=5 µm.

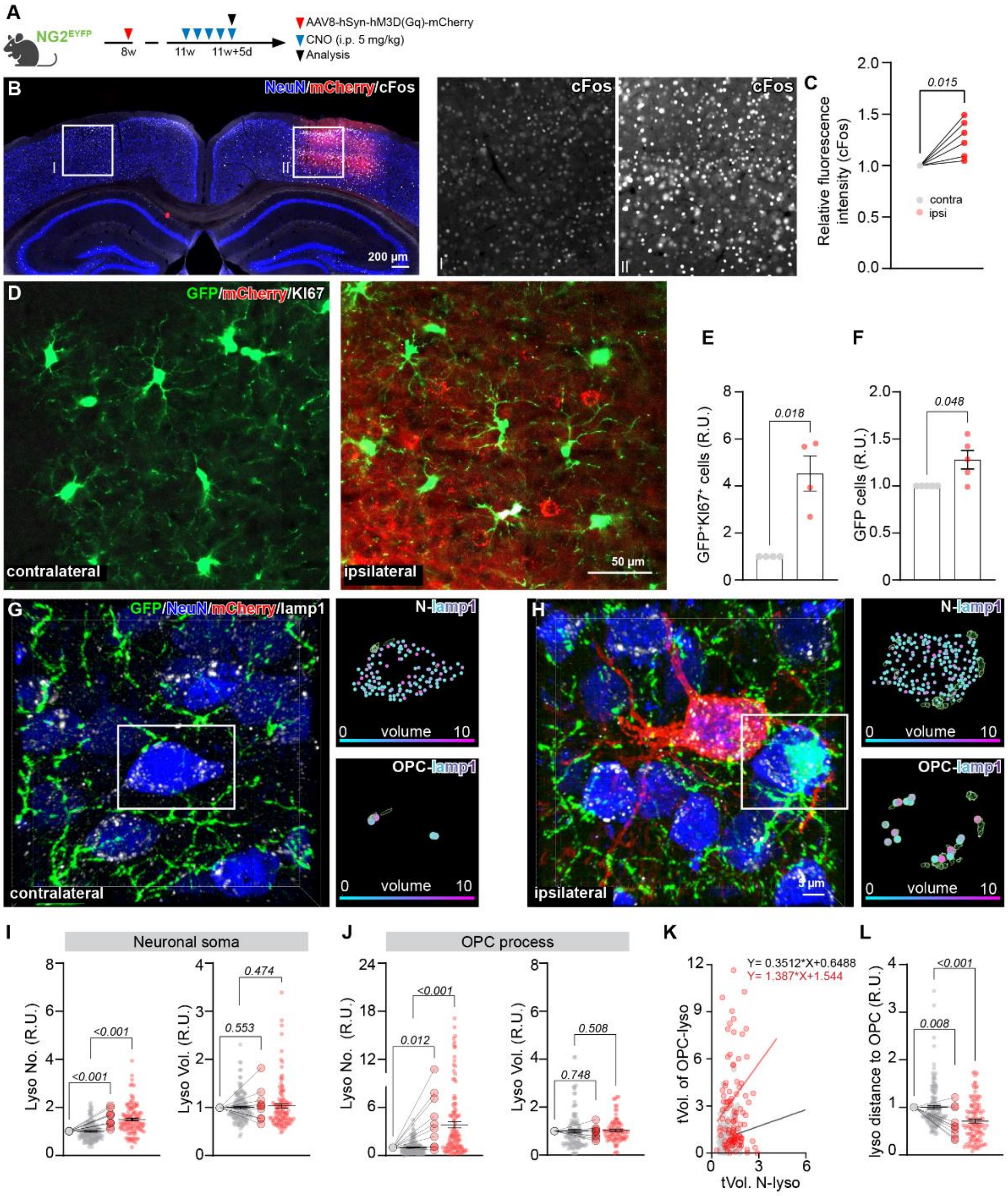


**Supplementary Figure 4. Chemogenetic activation of neurons triggers formation of OPC-neuron junction and increase of lysosome volume in neurons and OPCs. A** Scheme of experimental plan. **B** Overview of coronal brain slices from NG2$^{EYFP}$ mice, intracortically injected with AAV8-hSyn-hM3D(Gq)-mCherry virus, immunostained with NeuN and cFos. mCherry expression indicates neurons successfully transfected with the virus. **C** Quantification of cFos fluorescence intensity from contra- and ipsilateral cortex. **D** Immunostaining of OPCs with GFP and Ki67 at the contra- and ipsilateral side of virus injected NG2$^{EYFP}$ mouse cortex. **E, F** Quantification of OPC proliferation and density based on GFP and Ki67 expression. **G, H** Neurons, OPCs and lysosomes were immunolabeled with NeuN (blue), GFP (green) and Lamp1 (white), respectively, and mCherry labels hSyn-hM3Dq transfected neurons. **I, J** Quantification of relative umber and volume of lysosomes in neuronal soma (**I**) and OPC processes (**J**) in the contra- and ipsilateral side. (Neuronal soma: contra: n=145 cells and N=10 mice, ipsi: n=114 cells N=10 mice; OPC processes: contra: n=143 cells and N=10 mice, ipsi: n=115 cells N=10 mice, two-sided unpaired t-tests for the cells anlysis and two-sided paired t-tests for the mice). **K** Quantitative analysis of total volume of lysosomes in OPC processes (tVol. of OPC-lyso) and in neurons (tVol. of N-lyso) (contra: n=145 cells and N=10 mice, ipsi: n=107 cells and N=10 mice). **L** Quantification of relative distance between lysosomes and OPC surface (contra: n=146 cells and N=10 mice, ipsi: n=114 cells N=10 mice, two-sided unpaired t-tests for the cells anlysis and two-sided paired t-tests for the mice). Scale bars in **B**=200 μm, **D**=50 μm, **H**=5 μm.

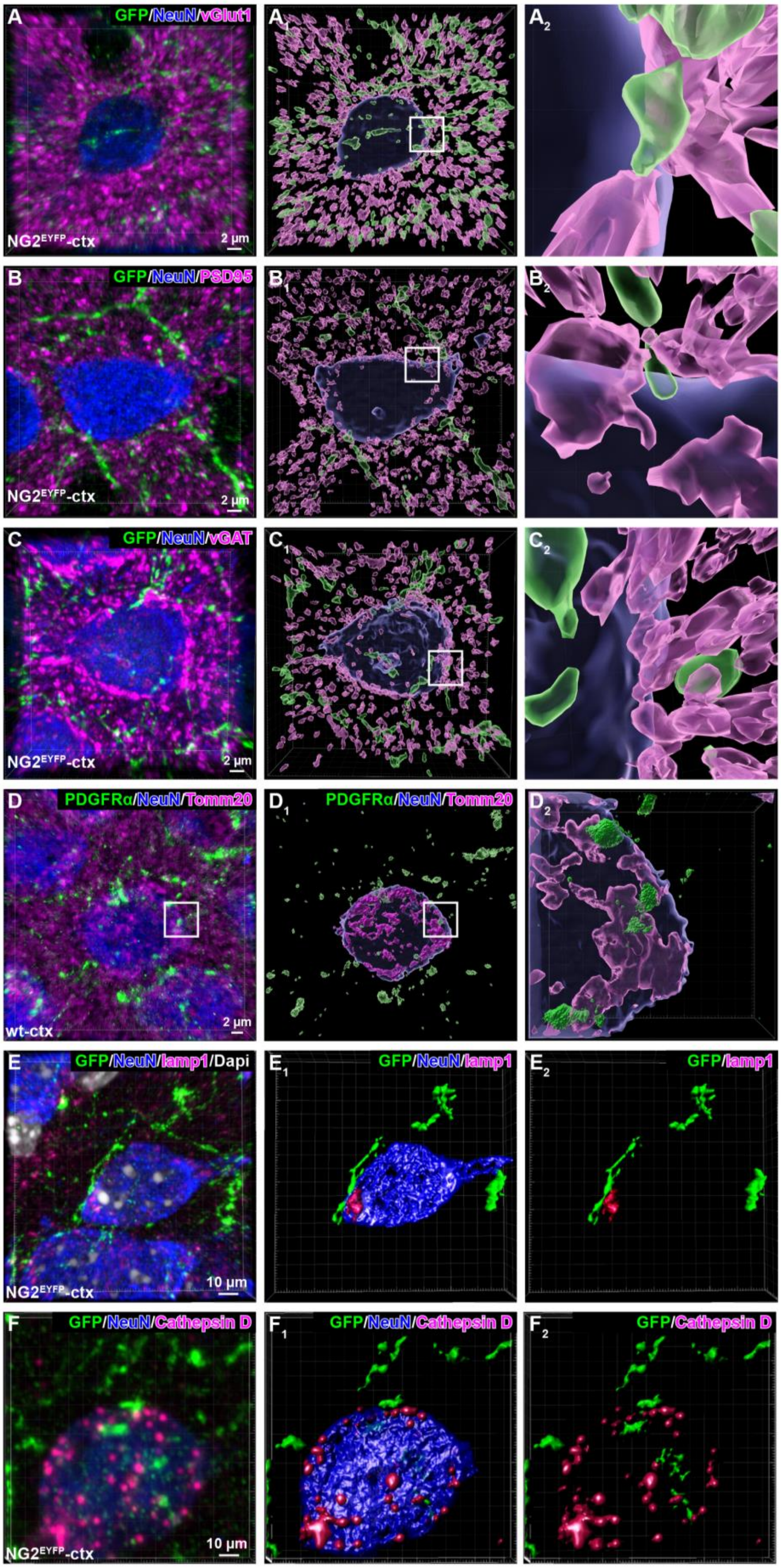

A GFP/NeuN/vGlut1
NG2EYFP-ctx
2 µm
A1
A2
B GFP/NeuN/PSD95
NG2EYFP-ctx
2 µm
B1
B2
C GFP/NeuN/vGAT
NG2EYFP-ctx
2 µm
C1
C2
D PDGFRα/NeuN/Tomm20
wt-ctx
2 µm
D1 PDGFRα/NeuN/Tomm20
D2
E GFP/NeuN/lamp1/Dapi
NG2EYFP-ctx
10 µm
E1 GFP/NeuN/lamp1
E2 GFP/lamp1
F GFP/NeuN/Cathepsin D
NG2EYFP-ctx
10 µm
F1 GFP/NeuN/Cathepsin D
F2 GFP/Cathepsin D

**Supplementary Figure 5. Neuronal lysosomes are present at the OPC-neuron contact site. A-C** Immunostaining of OPC and neurons with GFP and NeuN in $NG2^{EYFP}$ mice in combination with synaptic proteins (presynaptic vGlut or vGAT and postsynaptic PSD95). **D** Immunostaining of mitochondria in OPCs and neurons with Tomm20, PDGFRα and NeuN in wildtype brain, respectively. **E, F** Immunostaining of lysosomes in OPCs and neurons with Lamp1 (**E**) or Cathepsin D (CTSD, **F**), GFP and NeuN in $NG2^{EYFP}$ mouse cortex. Scale bar in **A-D**=2 µm, **E, F**=10 µm.

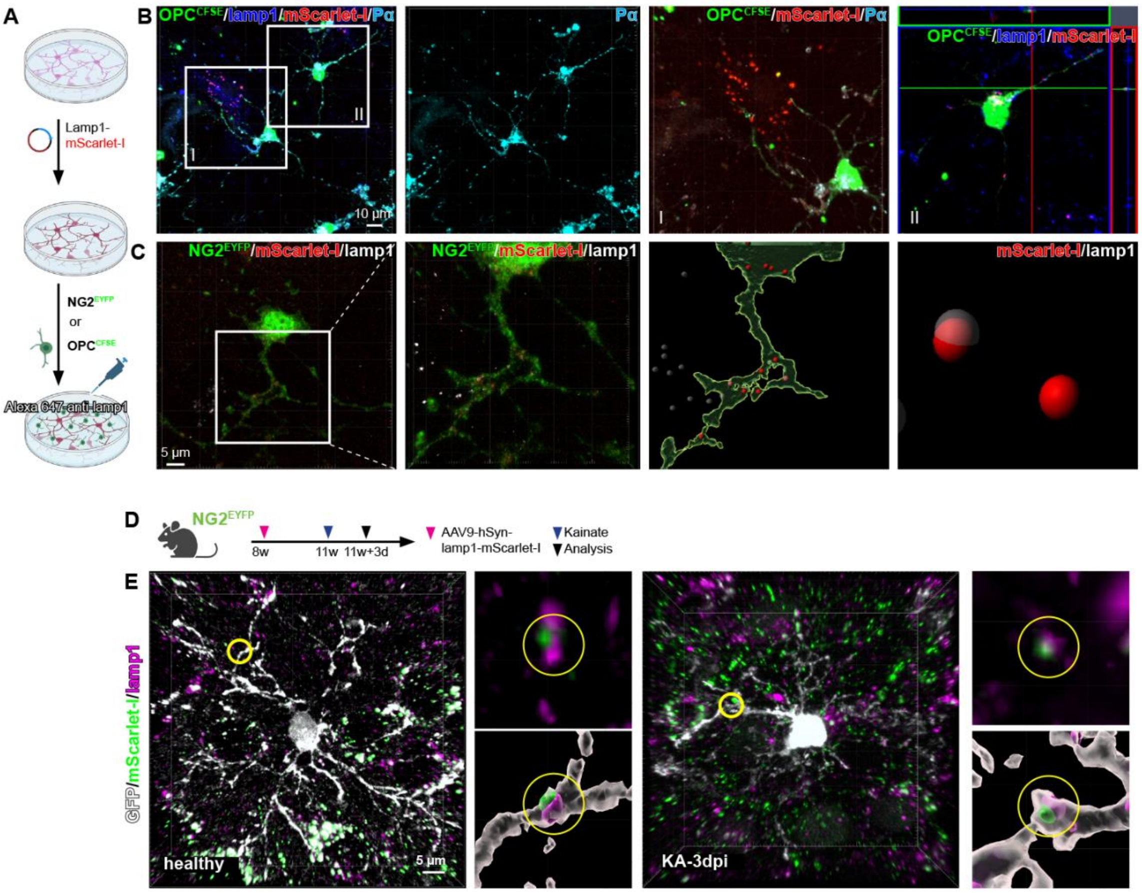


**Supplementary Figure 6. Neuronal lysosome trafficking and release is associated with OPC contact. A** Experimental scheme. mScarlet-I is expressed in neuronal lysosome. **B** Immunostaining of lysosomes with Lamp1 and OPCs were identified by CFSE-cell trace or EYFP combined with PDGFRα (Pα) expression in OPC-neuron co-culture. mScarlet-I expressing neuronal lysosomes appeared in OPCs, with some of them bound with DsRed-antibody targetting mScarlet added to the medium. **D** Experimental scheme of **E**. **E** Brain slices from healthy (left) and the kainate (KA) injected (right) mice were immunostained with GFP (OPC) and Lamp1 l(yososome). mScarlet-I is initially expressed by neuronal lysosomes. Three-dimensional reconstruction of the image shows mScarlet-I-expressing lysosomes in OPC processes. Scale bar in **A**=10 µm, in **C, E**=5 µm.

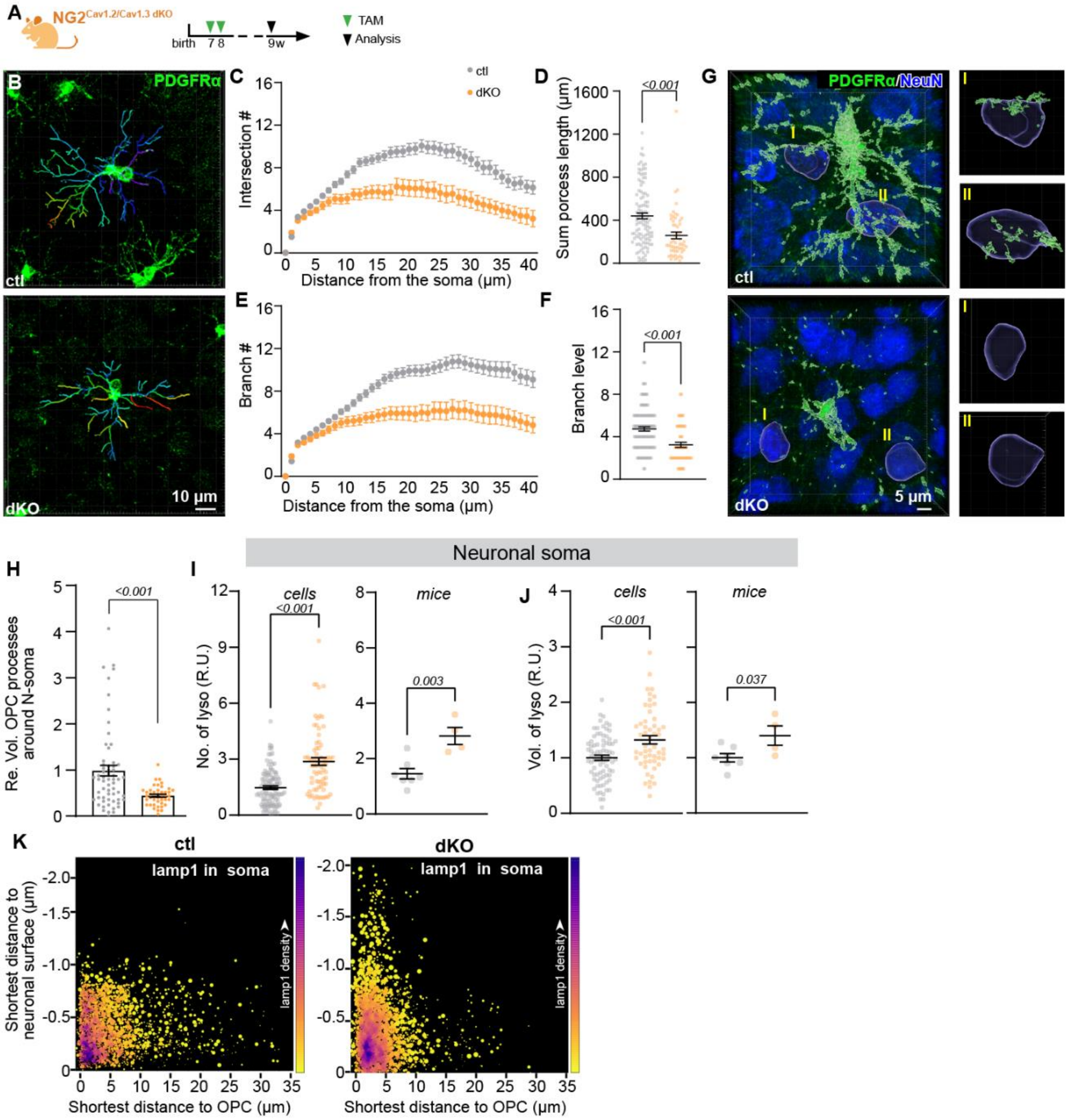


**Supplementary Figure 7. Neuronal lysosomes are present at the OPC-neuron contact sites.** **A** Experimental scheme for conditional deletion of Cav1.2 and Cav1.3 in OPCs. **B** Exemplary images of OPCs stained with PDGFRα and 3D reconstructed with Imaris. **C-F** Morphological analysis of OPCs from control (ctl) and dKO mouse cortex. **C** Mean distribution plots of the number of sholl intersections as a function of the distance from the OPC soma. **D** Comparison of total length of OPC processes between ctl and dKO groups. **E** Distribution plot of branching levels of OPC process as a function of the distance from the OPC soma. **F** Comparison of mean branching levels of ctl and dKO OPCs. **G** Immunostaining and 3D reconstruction of OPCs and neurons with PDGFRα and NeuN in the cortex of ctl and dKO mice. **H I, J** Quantification of relative number and volume of lysosomes in all neurons of ctl and dKO mouse brain. **K** Quantitative analysis of lysosome volume and shortest distance to OPC surface and neuronal surface. The relative volume of each lysosome is indicated by the size of the circle and the density of the lysosomes were indicated by the colour code, with the purple indicating higher density. Scale bars in **A**=10 µm and **G**=5 µm.

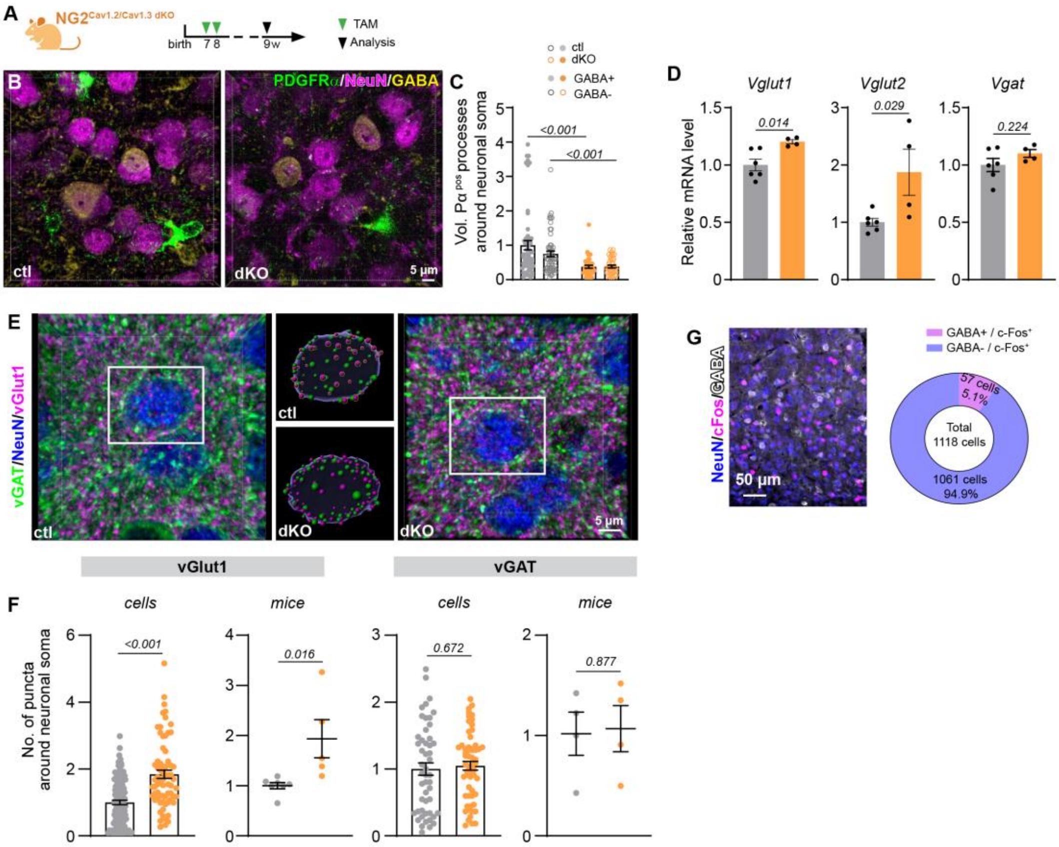

**Supplementary Figure 8. Reduced OPC-neuron contact increases excitatory but not inhibitory circuits. A** Experimental scheme. **B, C** The relative volume of OPC contact on GABAergic and non-GABAergic neurons was analysed after immunostaining of OPCs (PDGFRα, green), neurons (NeuN) and GABA (yellow). **D** Quantitative analysis of mRNA levels of vGlut1, vGlut2 and vGAT in the cortex of control and dKO mice. **E** Immunostaining of vGlut1 (magenta), NeuN (blue) and vGAT (green) in the cortex of ctl and dKO mice. **F** Quantification of vGlut1$^{+}$ or vGAT$^{+}$ puncta on neuronal somata in control and dKO mouse cortex. **G** Immunostaining of NeuN, cFos and GABA in wildtype mouse cortex shows that majority of cFos$^{+}$ neurons are non-GABAergic neurons. Scale bar in **B, E**=5 µm, **G**=50 µm.

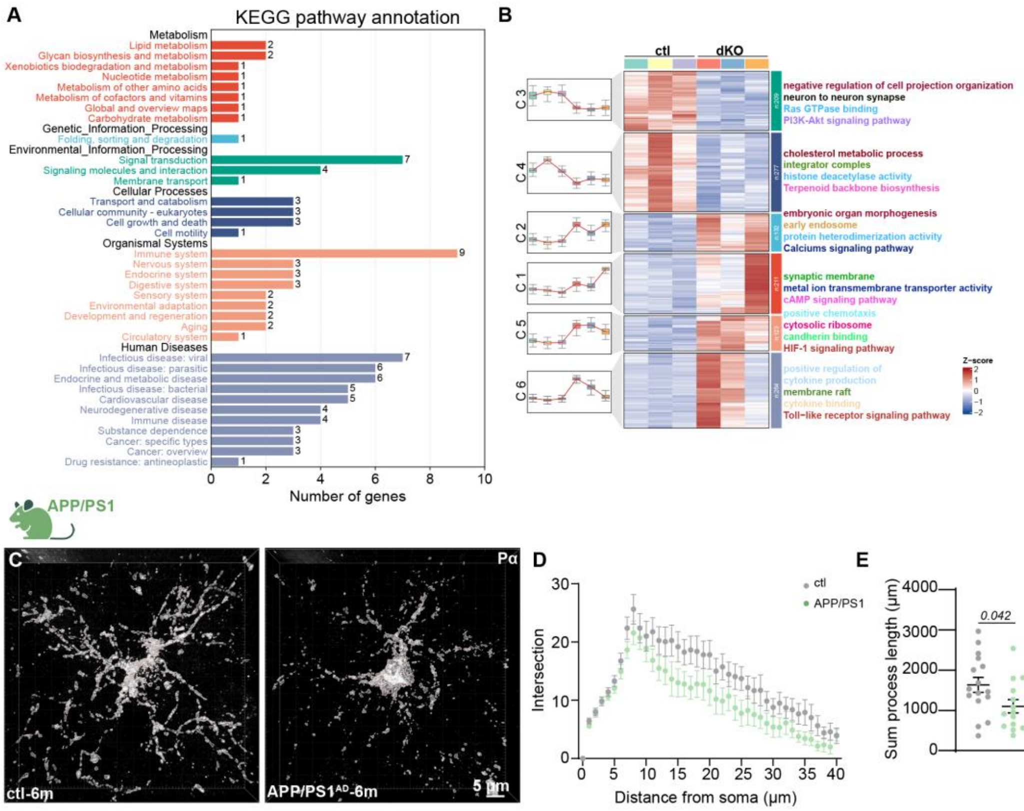


**Supplementary Figure 9. Reduced OPC arborization is associated with neurodegeneration. A, B** KEGG and biological processes analysis of genes altered in the dKO OPCs. **C** Three-dimensional reconstructions of OPCs based on their PDGFRα (Pα) immunoreactivity from the 6 month old APP/PS1 AD mouse model and age-matched control mouse cortex. **D, E** Morphological analysis of OPC processes for their number of intersections and sum length. Scale bar=5 μm.
